# Scaling Molecular Representation Learning with Hierarchical Mixture-of-Experts

**DOI:** 10.1101/2025.08.09.669511

**Authors:** Xiang Zhang, Shenbao Yu, Jie Xia, Fan Yang

## Abstract

Recent advancements in large-scale self-supervised pretraining have significantly improved molecular representation learning, yet challenges persist, particularly when addressing distributional shifts (e.g., under scaffold-split). Drawing inspiration from the success of Mixture-of-Experts (MoE) networks in NLP, we introduce H-MoE, a hierarchical MoE model tailored for molecular representation learning. Since conventional routing strategies struggle to capture global molecular information—such as scaffold structures, which are crucial for enhancing generalization—we propose a hierarchical routing mechanism. This mechanism first utilizes scaffold-level structural guidance before refining molecular characteristics at the atomic level. To optimize expert assignment, we incorporate scaffold routing contrastive loss, ensuring scaffold-consistent routing while preserving discriminability across molecular categories. Furthermore, a curriculum learning approach and dynamic expert allocation strategy are employed to enhance adaptability. Extensive experiments on molecular property prediction tasks demonstrate the effectiveness of our method in capturing molecular diversity and improving generalization across different tasks.

## 1 Introduction

Accurate and efficient molecular representation is a critical task in drug property prediction[1]. Traditional methods primarily rely on handcrafted molecular descriptors combined with machine learning models, such as regression models based on random forests[2] or support vector machines[3]. While these approaches exhibit stable performance on specific tasks, their static molecular representations struggle to capture the complex global and contextual information inherent in molecular structures. Additionally, they are constrained by dataset scale and feature engineering quality[4].

In recent years, the rise of large-scale self-supervised pretraining models has fundamentally transformed traditional molecular feature learning paradigms[5][6][7]. Several notable molecular pretraining models, such as Graphormer[8] and Mole-BERT[9], have demonstrated superior performance in molecular property prediction and drug discovery by learning rich molecular representations from vast unlabeled datasets[10]. These models transcend the limitations of manually crafted features and improve adaptability and transferability through extensive data-driven learning.

Despite these advances, current methods still exhibit notable weaknesses in handling molecular diversity and task complexity, particularly under distributional shifts (such as using scaffold-split[11]) where model performance significantly deteriorates[12][9]. Mixture of Experts (MoE) networks have achieved remarkable success in the Natural Language Processing (NLP) domain, emerging as a key strategy in state-of-the-art Large Language Models (LLMs)[13][14][15][16]. MoE’s core principle—expert specialization—enhances generalization across various tasks, while its sparse activation mechanism ensures efficient knowledge transfer[17][18][19]. Given the important role of generalization in molecular representation learning, we introduce the first MoE-based framework tailored for molecular representations, along with several key innovations designed to address existing challenges.

**Firstly**, directly using atomic-level routing strategies (e.g., routing based on individual atoms) are insufficient for capturing global structural features in drug-like molecules, limiting model generalization in complex molecular systems. The global features of a molecule, such as its scaffold, are closely intertwined with the properties it exhibits[12]. To overcome this, we propose a **hierarchical scaffold-atom routing** mechanism that first routes based on the global molecular scaffold, followed by refined routing using local atomic features. This approach enables different experts to specialize in distinct scaffold categories—such as aromatic compounds, aliphatic chain molecules, and heterocyclic systems—enhancing the model’s ability to represent structural diversity (Figure1).

**Figure 1:**
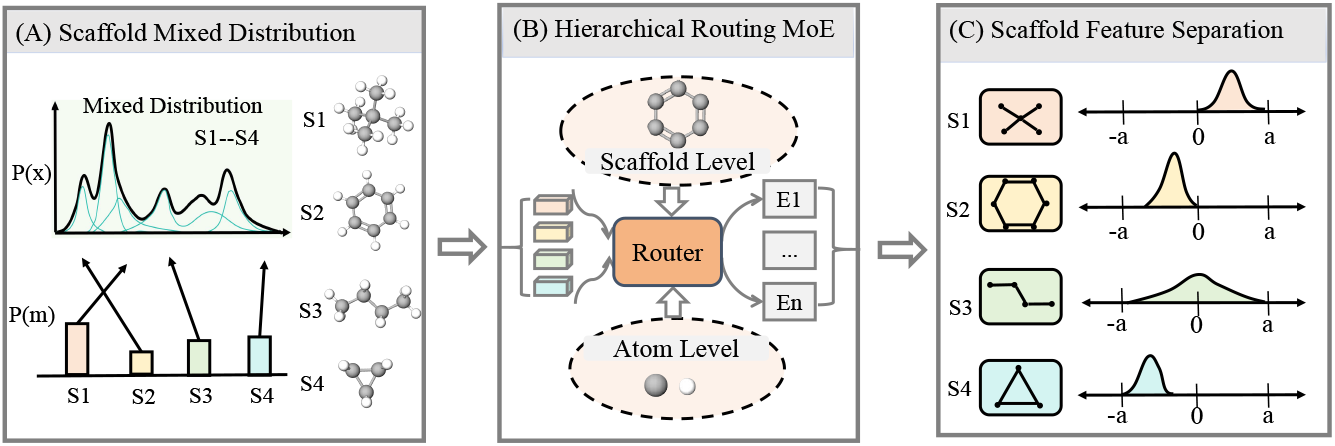
The left side (A) illustrates the overall distribution of diverse molecular scaffolds in the pretraining dataset, exhibiting a mixed probabilistic distribution. Through the central routing (B), which integrates both scaffold-level and atom-level assignments, molecules are automatically directed to different expert networks. This process ultimately achieves the scaffold feature separation depicted on the right side (C).

**Secondly**, to further optimize scaffold-level expert assignment, we introduce a **scaffold routing contrastive loss**, leveraging contrastive learning to refine scaffold-level routing decisions. Specifically, molecules sharing the same scaffold should exhibit similar routing distributions, whereas those belonging to different scaffolds must be distinctly separated. This loss function mitigates redundant learning across experts and enhances routing discriminability.

**Thirdly**, to ensure stable scaffold-level routing while adapting to molecular diversity, we introduce a **curriculum learning strategy** alongside a **dynamic expert allocation** mechanism that adjusts the number of activated experts based on scaffold distribution. Experimental results across multiple benchmarks demonstrate that our approach significantly improves molecular diversity modeling while maintaining computational efficiency.

**Scalability**. H-MoE is designed to efficiently process molecular data across various scales and structures, leveraging a scaffold-guided routing mechanism that dynamically adjusts expert activation based on scaffold distribution. Pre-training on 1.45 million molecules from ChEMBL[10] underscores its ability to handle large-scale data, while extensive evaluations on MoleculeNet[11] benchmark, covering classification and regression tasks, further validate its scalability. Additionally, the theoretical analysis in Appendix A establishes a solid foundation, demonstrating its capacity to generalize under distribution shifts and accommodate diverse data scales.

Our work presents several key contributions:

- We present a novel hierarchical MoE-based molecular representation learning framework, incorporating scaffold-guided routing to enhance atomic-level expert decisions. Additionally, we provide theoretical guarantees demonstrating the superiority of scaffold-guided MoE over standard MoE.
- We incorporate scaffold routing contrastive loss and an adaptive dynamic expert allocation strategy to refine expert assignment, enhancing generalization across diverse molecular structures.
- Our approach surpasses classical methods, including some using 3D conformations, by effectively leveraging stable 2D molecular features and an innovative routing mechanism to capture key characteristics.

## 2 Related work

### Molecular pretraining models

Molecular property prediction has seen extensive development in pretraining strategies[4], which can be grouped into three main methods. Contrastive learning methods, such as GraphCL[20] and AD-GCL[6], enhance molecular structure representation by distinguishing variations across samples. Masked reconstruction methods, exemplified by GraphMAE[7] and AttrMask[5], leverage masking mechanisms to capture local structural patterns. Multi-dimensional representation methods, like GraphMVP[21] and MOLEBLEND[22], integrate both 2D and 3D molecular features to improve structural completeness. While these methods advance molecular representation, challenges remain in handling the diversity and complexity of molecular data[12], highlighting the need for further refinement.

### Mixture of experts

In the NLP domain, Sparsely-Gated MoE[23] first introduced the mixture-of-experts paradigm, laying a strong theoretical foundation for efficient training. Subsequent advancements[24][16][15] introduced several key improvements, enabling the effective pretraining of large-scale foundational models such as LLMs (e.g., DeepSeek[13]). Various studies [25][19][26]explored different expert-assigment strategies in MoE models and compared their performance. Additionally, some works[13][14][27][17]focused on optimizing the expert networks within MoE to improve overall model utilization. Furthermore, significant advancements[18][28] were made in enhancing the operational efficiency of MoE models. Inspired by the success of MoE in NLP, we propose a hierarchical mixture-of-experts network designed specifically for molecular.

### Molecular scaffold

A molecular scaffold is the core structure of a molecule, playing a fundamental role in drug design [29]. Studies have shown [30] that scaffolds significantly influence chemical and biological activity, while recent research[31] has explored scaffold modification strategies for drug optimization. Scaffold split[11], a widely used dataset partitioning method, ensures distinct scaffolds in training and test sets, offering a more rigorous evaluation of model generalization[32] compared to random splitting, which may lead to overly optimistic performance estimates. Our approach incorporates scaffolds as the first-level routing mechanism in the MoE model, effectively improving its ability to generalize across scaffold domains in real-world drug discovery applications.

## 3 Preliminary

The molecular graph is represented as 𝒢 = (𝒱, *ε*), where 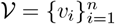 denotes the set of atoms, and ε = {(𝓋_*i*_, 𝓋_*j*_, *𝓇*_*ij*_)} represents chemical bonds with bond types 𝓇_*ij*_. The initial atomic features are generated via a learnable embedding layer 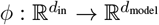.

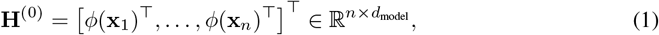

where **x**_*i*_ contains atomic properties such as atom type and formal charge.

The model consists of *L* stacked Transformer layers, each containing multi-head attention (MHA), feed-forward networks (FFN), and layer normalization (LayerNorm). The attention mechanism first projects input features using learnable matrices:

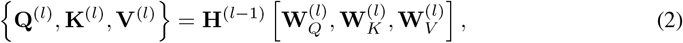

followed by multi-head attention computation:

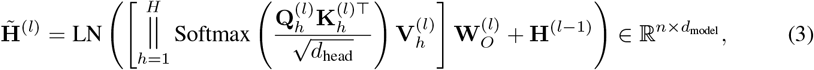

where *∥* denotes head-wise concatenation, and 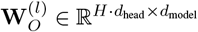 is the output projection matrix.

The features are further processed by a feed-forward network (FFN), which acts as an expert in our HMoE model.

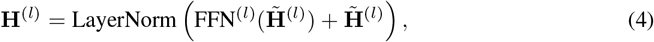

The global molecular representation is obtained via a [CLS] token and projected to property predictions: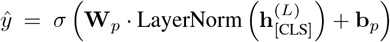, where *σ* is an activation function, 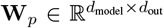 and 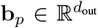 are projection parameters, forming an end-to-end Transformer-based molecular property prediction framework.

## 4 Method

### 4.1 Atom-level routing

Given a molecular sequence *S* = *{𝓋*_1_, *𝓋*_2_, …, *𝓋*_*n*_*}*, where each atom *v*_*i*_ *∈ 𝒱* is associated with a hidden state 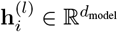 at layer *l*, we introduce a atom-level routing mechanism that dynamically assigns each atom to a subset of *k* experts.

The routing module computes a gating distribution **G**^(*l*)^ *∈ ℝ*^*n×N*^ for all atoms, where *N* is the number of experts, via a learnable routing transformation:

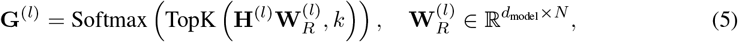

where: TopK(•, *k*) retains only the top-*k* expert assignments per atom, enforcing sparsity.

Each expert 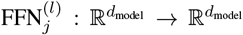 processes its assigned atoms independently. The expert-aggregated output for atom *v*_*i*_ is computed as:

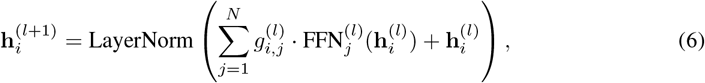

where 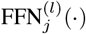 applies a two-layer MLP with ReLU activation and expert-specific parameters, the gating score 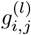 represents the probability of atom 𝓋_*i*_ being routed to expert 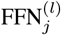.

### 4.2 Scaffold-level routing

#### Algorithm 1

Scaffold-Level Expert Router by Contrastive Learning

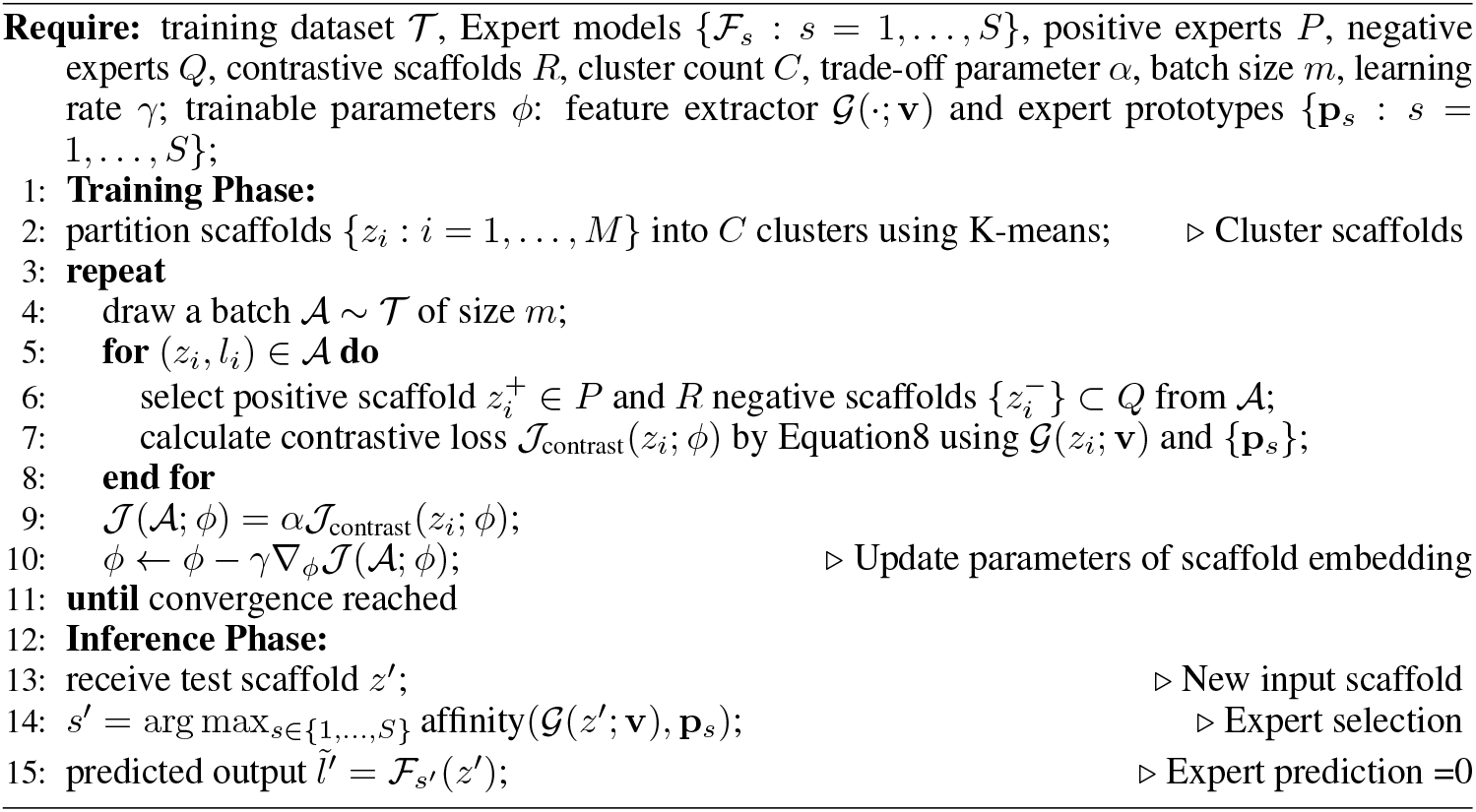

#### Motivation

Unlike natural language, local and global information of chemical molecules shows significant differences. Different types of substructures or molecular scaffolds are likely to correspond to distinct chemical properties, such as solubility and toxicity. Therefore, the input here comprehensively considers both atom-level and molecular scaffold-level feature. Guided by the prior knowledge of molecular scaffold features, a hierarchical routing strategy with a two-level router network is designed (Figure 2).

**Figure 2:**
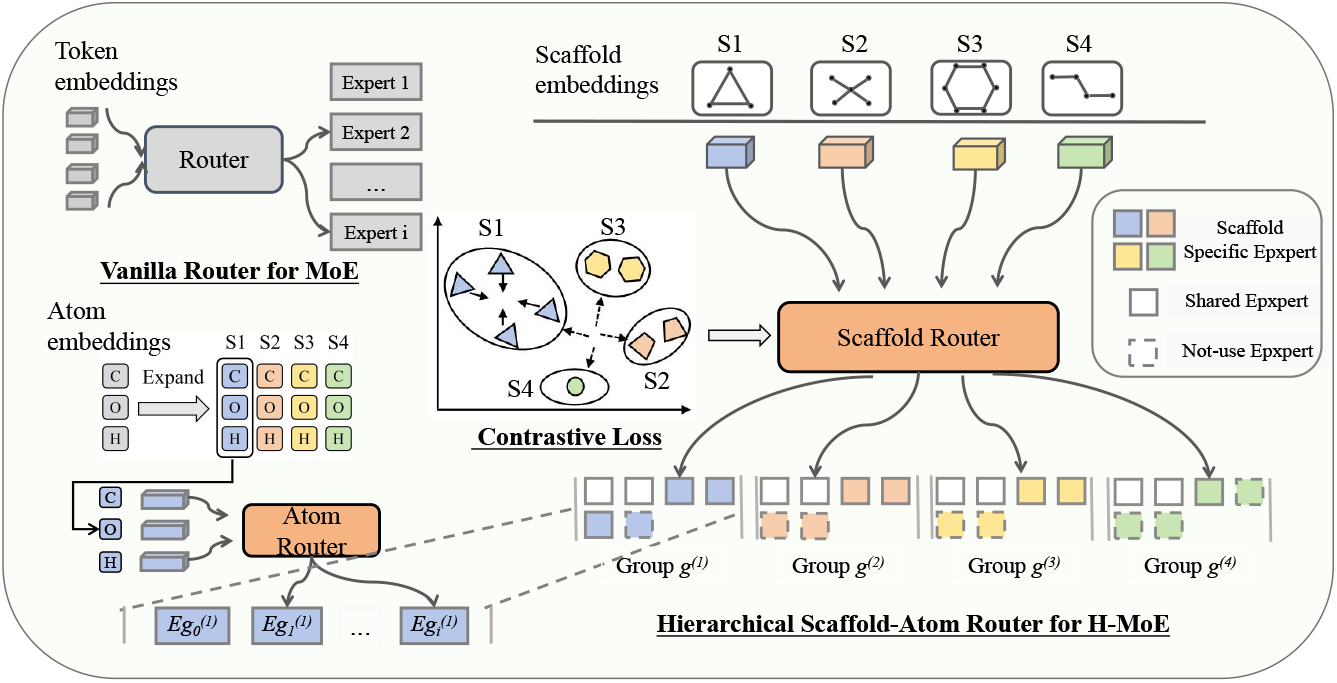
The hierarchical MoE architecture for molecular representation consists of two core components. Firstly, the Scaffold Router partitions input molecules based on their scaffold structures into distinct groups (Group g^(1)^ to Group g^(4)^), with each group corresponding to a class of structurally similar compounds. Within this framework, Scaffold-Specific Experts are responsible for maintaining local consistency within a specific scaffold feature distribution, while Shared Experts (white) capture general chemical rules across different scaffold categories. The number of experts assigned to each scaffold category varies based on scaffold diversity. Secondly, the Atom Router further refines routing within each scaffold group at the atomic level (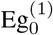 to 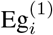), enhancing the model’s ability to capture local chemical environments with greater precision.

#### Scaffold-level routing

Specifically, for each input molecule (denoted as *x*): (1) The Morgan fingerprint vector *F*_*morgan*_ = *ϕ*(*f*_*i*_ *• r*_*i*_) of the target molecular scaffold is extracted, where *f*_*i*_ represents atomic features, *r*_*i*_ denotes the connection relationship between atoms, and *ϕ* is the fingerprint generation function. Then, a simple fully connected layer *W* generates the scaffold embedding *E*_*scaffold*_ = *σ*(*W • F*_*morgan*_ + *b*). (2) The scaffold router takes the molecular scaffold embedding *E*_*scaffold*_ as input and outputs the probability distribution of experts from the perspective of molecular scaffolds 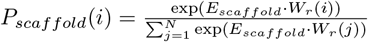, where *W*_*r*_(*i*) is the weight parameter associated with the *i*-th candidate expert, and *N* is the total number of candidate experts. Then, the Top *k* function is applied to reduce the candidate experts to guide atom-level routing.

##### Theorem 1

(Scaffold-Aware Routing Benefit). *Our scaffold-aware routing mechanism guarantees superior generalization compared to standard MoE architectures. Formally, under scaffold-partitioned data distribution (Def. A*.*1), the MoE model with scaffold-aware routing achieves:*

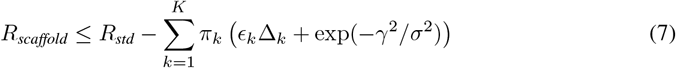

*where R*_*scaffold*_ *and R*_*std*_ *are the expected risks of scaffold-aware and standard* 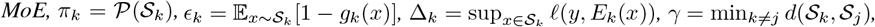 *and σ is the Lipschitz constant of h*_*θ*_. *(Proof: Appendix A)*

#### Scaffold routing contrastive loss

Due to the instability of the original scaffold features, a staged training strategy is adopted: in the first stage, a molecular scaffold embedding layer is pre-trained by contrastive learning to make the expert selection behavior of different scaffolds more specific; in the second stage, the scaffold embedding layer is shared, and molecular masking pre-training is performed. Based on the original clustering scaffold labels *y*, inter-scaffold contrastive learning is implemented to align the expert probability distributions of scaffolds with the same label while distancing different scaffolds. The scaffold router contrastive loss *L*_contrast_ is defined as :

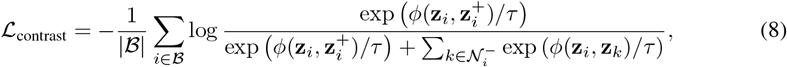

where *ϕ*(•, *•*) denotes the cosine similarity between normalized scaffold embeddings **z**, ℬ is the mini-batch, 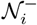 represents negative samples for anchor *i*, and *τ* controls the softmax temperature.

The scaffold routing behavior is optimized through contrastive learning, which drives the collaborative optimization of expert embeddings and molecular scaffold features. The process is divided into training and inference stages, see Algorithm 1 for details.

### 4.3 Curriculum learning and dynamic expert allocation

#### Curriculum learning

During the pretraining phase, to ensure that the scaffold-level routing strategy more stably guides the atom-level routing process, we adopt a curriculum learning strategy. Specifically, we define the weighted cosine similarity Sim between the atom-level expert selection probability distributions of two consecutive steps as the criterion for training stability. The formula is as follows:

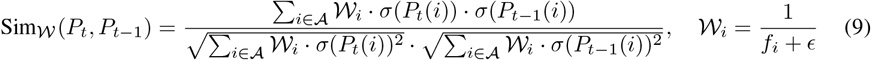

where *σ*(*P*_*t*_(*i*)) and *σ*(*P*_*t* − 1_(*i*)) represent the expert selection probability of atom type *i* at the current step and the previous step, respectively. The weight 𝒲_*i*_ is defined as the inverse of the occurrence frequency *f*_*i*_ of atom type *i*, with *ϵ* as a smoothing factor to prevent numerical instability when the frequency is zero.

#### Dynamic expert allocation

To address scaffold clustering imbalance and expert training instability, we implement a dynamic allocation strategy. Scaffolds are grouped into high, medium, and low subsets based on sample count. For each subset *S*_*j*_, expert number *k*_*l,j*_ is iteratively adjusted using validation metric 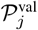 and threshold *λ*. If 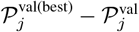 is below *λ, k*_*l,j*_ increases; otherwise, it decreases, with further refinement over Δ*n* iterations. Once validation stabilizes, *k*_*l,j*_ is fixed, finalizing training and determining the scaffold-specific routing top-*k*_*l,j*_. The dynamic expert allocation process is outlined in Algorithm 2 of Appendix B.

During molecular processing, the random initialization of expert FFN layers introduces uncertainty, causing imbalances where certain atom subsets over-rely on a few experts or experts specialize in limited atoms. This leads to extreme load imbalance, weakening the H-MoE layer’s effectiveness. To mitigate this, we introduce a balancing loss function *L*_*bl*_, defined as follows:

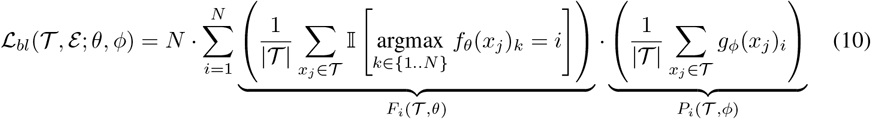

where 𝒢 denotes the atom batch with cardinality ∣ 𝒢 *∣, θ* and *ϕ* parameterize the routing and probability systems respectively, and 𝕀 [•] is the indicator function. Two key components: *F*_*i*_(𝒢, *θ*) represents atom routing frequency of expert *i* via hard assignments; *P*_*i*_(𝒢, *ϕ*) represents atom mean activation probability for expert *i* under soft routing.

## 5 Experiments

### 5.1 Experimental setup

#### Dataset

For pretraining, we utilize 1.45 million molecules from ChEMBL[10] for masked pretraining[33]. For fine-tuning, we employ eight benchmark classification datasets from MoleculeNet[11], such as MUV[34], Bace[35], BBBp[36], Sider[37], ClinTox[38]. Additionally, we conducted experiments on five commonly used regression datasets, including ESOL[39], FreeSolv[40], Lipo[41], Malaria[42], and CEP[43]. To ensure a robust evaluation of model performance across varying data distributions, we adopt the scaffold-based train/test split method, following the approach used in [9]. A more detailed description of the datasets is provided in AppendixC.1.

#### Baseline

We select several classic methods in molecular property prediction as benchmarks. The first category consists of contrastive learning-based approaches that enhance model performance by comparing differences between samples, including EdgePred[44], ContextPred[5], GraphLoG[45], G-Contextual[46], G-Motif[46], AD-GCL[6], JOAO[47], SimGRACE[48], and GraphCL[20]. The second category comprises self-supervised masked and reconstruction-based learning methods, which extract molecular features through masking or reconstruction, such as GraphMAE[7], MGSSL[49], AttrMask[5], Mole-BERT[9], StructMAE[50], and Hi-GMAE[51]. The final category includes molecular representation methods that integrate 3D molecular information to improve performance, encompassing 3D InfoMax[52], GraphMVP[21], Unimol[53], MoleculeSDE[54], and MOLEBLEND[22]. Since 3D-based approaches incorporate additional structural information compared to the other two categories, we list them separately.

#### Experimental setup

For backbone model, we adopt a transformer model with 6 layers, each containing 4 attention heads, and set the embedding dimension to 256. Following [55], we apply attention masking to leverage the 2D graph information of molecules. The pretraining process utilizes the Adam optimizer with a learning rate of 1e-4, training for 20 epochs. The masking ratio for pretraining is set at 0.15, ensuring that molecules with very few atoms have at least one masked atom. For fine-tuning downstream tasks, we use a two-layer MLP architecture, with specific hyperparameter settings detailed in Appendix C.2. Also see Appendix C.3 for details on the division of scaffold.

### 5.2 Main experiments

We evaluate the performance of H-MoE on MoleculeNet, a widely used benchmark dataset encompassing various tasks such as drug discovery, quantum chemistry, and biological activity prediction. As shown in the Table 1, H-MoE achieves superior average performance compared to all prior methods, outperforming existing models on six out of eight datasets, even when trained using only 2D information and masked pretraining strategies. The only exceptions are the BBBP and MUV datasets, where H-MoE falls short of 3D-based methods such as MolBlend and MoleculeSDE. On some datasets, H-MoE demonstrates significant improvements over previous approaches (e.g., 90.6 vs. 87.9 on ClinTox). Additionally, in regression tasks of Table 2, H-MoE achieves the best results on three out of five datasets and ranks second on the remaining two. These strong results validate the effectiveness of the H-MoE, even in scenarios with scaffold domain shift.

**Table 1:**
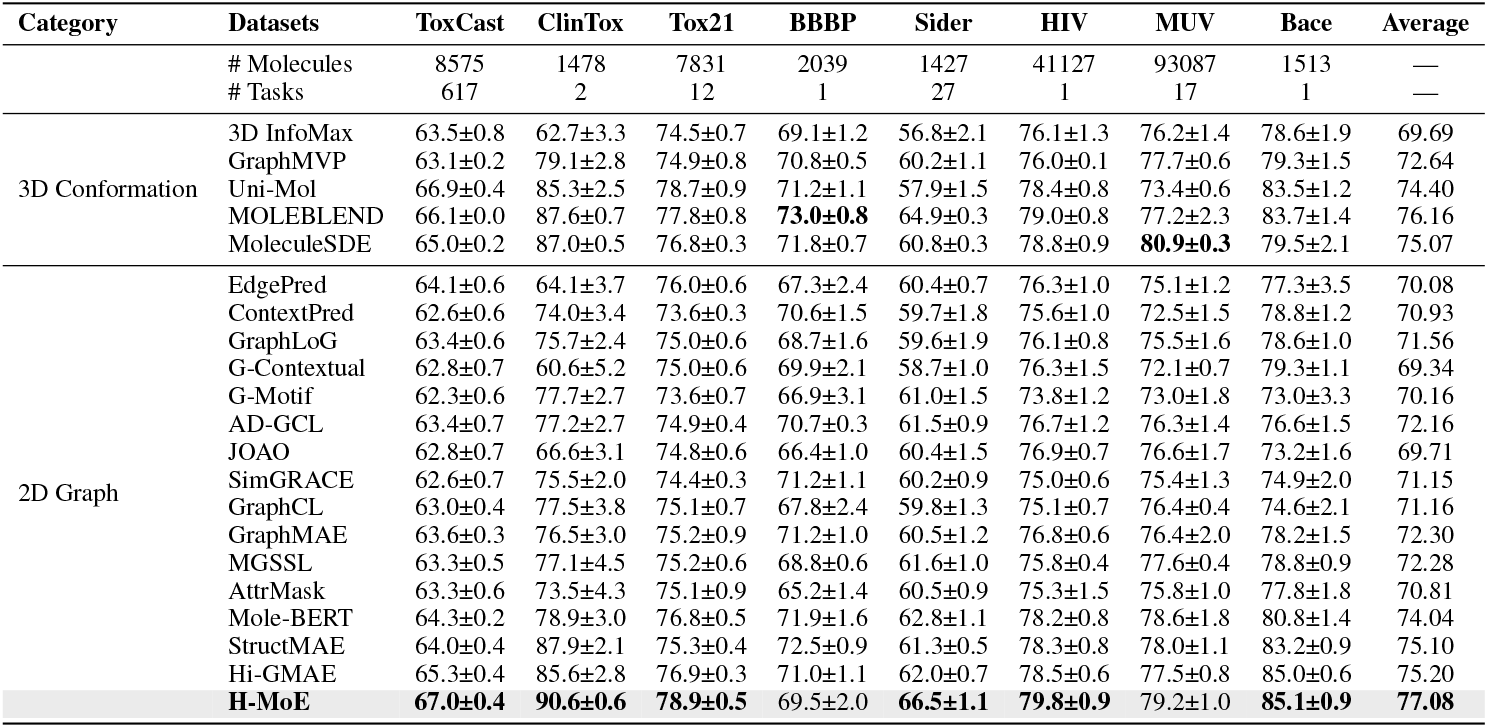
Results on molecular property classification tasks (ROC-AUC %, higher is better ↑). **Bold** indicates the best performance.

| Category | Datasets | ToxCast | ClinTox | Tox21 | BBBP | Sider | HIV | MUV | Bace | Average |
| --- | --- | --- | --- | --- | --- | --- | --- | --- | --- | --- |
|  | # Molecules | 8575 | 1478 | 7831 | 2039 | 1427 | 41127 | 93087 | 1513 | — |
|  | # Tasks | 617 | 2 | 12 | 1 | 27 | 1 | 17 | 1 | — |
| 3D Conformation | 3D InfoMax | 63.5±0.8 | 62.7±3.3 | 74.5±0.7 | 69.1±1.2 | 56.8±2.1 | 76.1±1.3 | 76.2±1.4 | 78.6±1.9 | 69.69 |
| | GraphMVP | 63.1±0.2 | 79.1±2.8 | 74.9±0.8 | 70.8±0.5 | 60.2±1.1 | 76.0±0.1 | 77.7±0.6 | 79.3±1.5 | 72.64 |
| | Uni-Mol | 66.9±0.4 | 85.3±2.5 | 78.7±0.9 | 71.2±1.1 | 57.9±1.5 | 78.4±0.8 | 73.4±0.6 | 83.5±1.2 | 74.40 |
| | MOLEBLEND | 66.1±0.0 | 87.6±0.7 | 77.8±0.8 | **73.0±0.8** | 64.9±0.3 | 79.0±0.8 | 77.2±2.3 | 83.7±1.4 | 76.16 |
| | MoleculeSDE | 65.0±0.2 | 87.0±0.5 | 76.8±0.3 | 71.8±0.7 | 60.8±0.3 | 78.8±0.9 | **80.9±0.3** | 79.5±2.1 | 75.07 |
| 2D Graph | EdgePred | 64.1±0.6 | 64.1±3.7 | 76.0±0.6 | 67.3±2.4 | 60.4±0.7 | 76.3±1.0 | 75.1±1.2 | 77.3±3.5 | 70.08 |
| | ContextPred | 62.6±0.6 | 74.0±3.4 | 73.6±0.3 | 70.6±1.5 | 59.7±1.8 | 75.6±1.0 | 72.5±1.5 | 78.8±1.2 | 70.93 |
| | GraphLoG | 63.4±0.6 | 75.7±2.4 | 75.0±0.6 | 68.7±1.6 | 59.6±1.9 | 76.1±0.8 | 75.5±1.6 | 78.6±1.0 | 71.56 |
| | G-Contextual | 62.8±0.7 | 60.6±5.2 | 75.0±0.6 | 69.9±2.1 | 58.7±1.0 | 76.3±1.5 | 72.1±0.7 | 79.3±1.1 | 69.34 |
| | G-Motif | 62.3±0.6 | 77.7±2.7 | 73.6±0.7 | 66.9±3.1 | 61.0±1.5 | 73.8±1.2 | 73.0±1.8 | 73.0±3.3 | 70.16 |
| | AD-GCL | 63.4±0.7 | 77.2±2.7 | 74.9±0.4 | 70.7±0.3 | 61.5±0.9 | 76.7±1.2 | 76.3±1.4 | 76.6±1.5 | 72.16 |
| | JOAO | 62.8±0.7 | 66.6±3.1 | 74.8±0.6 | 66.4±1.0 | 60.4±1.5 | 76.9±0.7 | 76.6±1.7 | 73.2±1.6 | 69.71 |
| | SimGRACE | 62.6±0.7 | 75.5±2.0 | 74.4±0.3 | 71.2±1.1 | 60.2±0.9 | 75.0±0.6 | 75.4±1.3 | 74.9±2.0 | 71.15 |
| | GraphCL | 63.0±0.4 | 77.5±3.8 | 75.1±0.7 | 67.8±2.4 | 59.8±1.3 | 75.1±0.7 | 76.4±0.4 | 74.6±2.1 | 71.16 |
| | GraphMAE | 63.6±0.3 | 76.5±3.0 | 75.2±0.9 | 71.2±1.0 | 60.5±1.2 | 76.8±0.6 | 76.4±2.0 | 78.2±1.5 | 72.30 |
| | MGSSL | 63.3±0.5 | 77.1±4.5 | 75.2±0.6 | 68.8±0.6 | 61.6±1.0 | 75.8±0.4 | 77.6±0.4 | 78.8±0.9 | 72.28 |
| | AttrMask | 63.3±0.6 | 73.5±4.3 | 75.1±0.9 | 65.2±1.4 | 60.5±0.9 | 75.3±1.5 | 75.8±1.0 | 77.8±1.8 | 70.81 |
| | Mole-BERT | 64.3±0.2 | 78.9±3.0 | 76.8±0.5 | 71.9±1.6 | 62.8±1.1 | 78.2±0.8 | 78.6±1.8 | 80.8±1.4 | 74.04 |
| | StructMAE | 64.0±0.4 | 87.9±2.1 | 75.3±0.4 | 72.5±0.9 | 61.3±0.5 | 78.3±0.8 | 78.0±1.1 | 83.2±0.9 | 75.10 |
| | Hi-GMAE | 65.3±0.4 | 85.6±2.8 | 76.9±0.3 | 71.0±1.1 | 62.0±0.7 | 78.5±0.6 | 77.5±0.8 | 85.0±0.6 | 75.20 |
| | **H-MoE** | **67.0±0.4** | **90.6±0.6** | **78.9±0.5** | 69.5±2.0 | **66.5±1.1** | **79.8±0.9** | 79.2±1.0 | **85.1±0.9** | **77.08** |

**Table 2:**
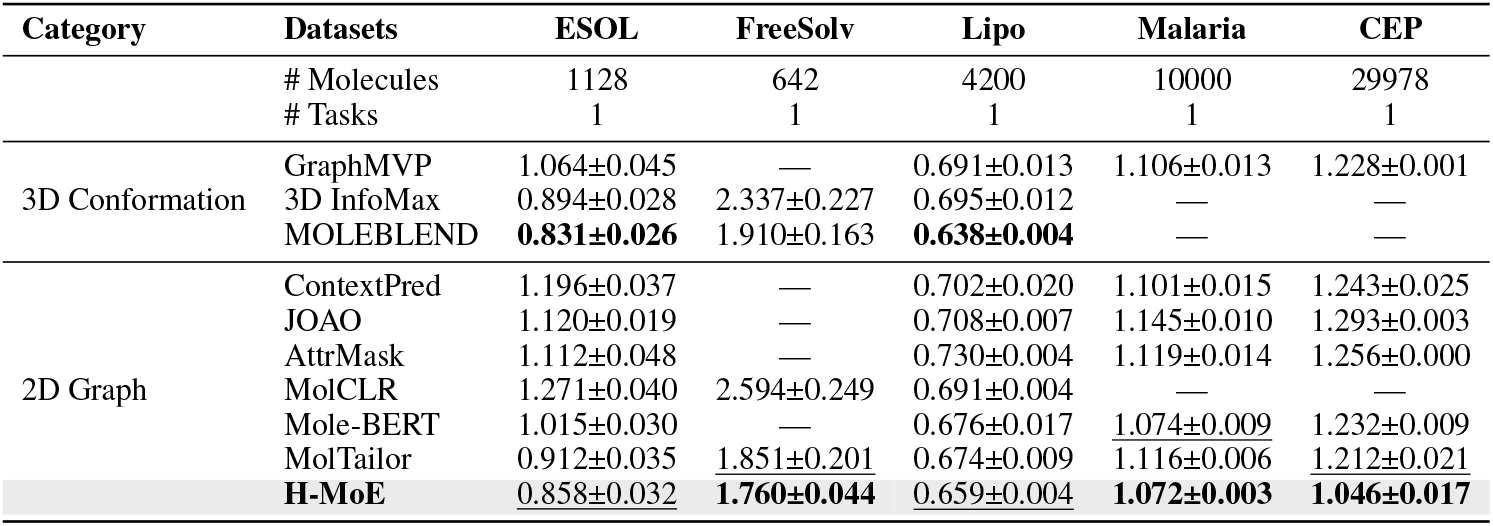
Results on molecular property regression tasks (RMSE %, lower is better ↓). **Bold** indicates the best performance and <u>underline</u> indicates the second best.

| Category | Datasets | ESOL | FreeSolv | Lipo | Malaria | CEP |
| --- | --- | --- | --- | --- | --- | --- |
|  | # Molecules | 1128 | 642 | 4200 | 10000 | 29978 |
|  | # Tasks | 1 | 1 | 1 | 1 | 1 |
| 3D Conformation | GraphMVP | 1.064±0.045 | — | 0.691±0.013 | 1.106±0.013 | 1.228±0.001 |
|  | 3D InfoMax | 0.894±0.028 | 2.337±0.227 | 0.695±0.012 | — | — |
|  | MOLEBLEND | **0.831±0.026** | 1.910±0.163 | **0.638±0.004** | — | — |
| 2D Graph | ContextPred | 1.196±0.037 | — | 0.702±0.020 | 1.101±0.015 | 1.243±0.025 |
|  | JOAO | 1.120±0.019 | — | 0.708±0.007 | 1.145±0.010 | 1.293±0.003 |
|  | AttrMask | 1.112±0.048 | — | 0.730±0.004 | 1.119±0.014 | 1.256±0.000 |
|  | MolCLR | 1.271±0.040 | 2.594±0.249 | 0.691±0.004 | — | — |
|  | Mole-BERT | 1.015±0.030 | — | 0.676±0.017 | <u>1.074±0.009</u> | 1.232±0.009 |
|  | MolTailor | 0.912±0.035 | <u>1.851±0.201</u> | 0.674±0.009 | 1.116±0.006 | 1.212±0.021 |
|  | **H-MoE** | <u>0.858±0.032</u> | **1.760±0.044** | <u>0.659±0.004</u> | **1.072±0.003** | **1.046±0.017** |

### 5.3 Ablation study

We conduct a comprehensive ablation study on eight classification datasets from MoleculeNet. As shown in the Table 3, H-MoE achieves optimal performance on five datasets, including ToxCast, ClinTox, and Sider, demonstrating the necessity of scaffold-level routing guidance as a global constraint for improving model performance across different scaffold distributions. Furthermore, even without scaffold-guided routing, adopting atomic-level routing as a local strategy still provides overall performance gains, outperforming densely structured baseline networks on multiple datasets. These results confirm the broad effectiveness of sparse routing strategies and their crucial role in enhancing model generalization capabilities.

**Table 3:** H-MoE ablation experiment. A: Atom-Level Routing. S: Scaffold-Level Routing.

| Method | ToxCast | ClinTox | Tox21 | BBBP | Sider | HIV | MUV | Bace |
| --- | --- | --- | --- | --- | --- | --- | --- | --- |
| w/o A, S | 65.7±0.5 | 88.6±0.9 | 77.8±0.9 | **70.1±2.5** | 64.1±1.2 | <u>78.5±1.3</u> | 76.3±1.5 | 84.2±1.1 |
| w/o S | <u>66.4±0.8</u> | 89.7±0.4 | **79.1±0.6** | 68.8±1.4 | <u>65.3±0.7</u> | 77.6±0.8 | 78.1±0.8 | **85.3±1.4** |
| H-MoE | **67.0±0.4** | **90.6±0.6** | <u>78.9±0.5</u> | <u>69.5±2.0</u> | **66.5±1.1** | **79.8±0.9** | **79.2±1.0** | <u>85.1±0.9</u> |

### 5.4 Other results

#### The role of scaffold-level routing contrastive loss

Figure 3a illustrates the cosine similarity matrix of scaffold-level router embeddings without applying scaffold contrastive loss, where off-diagonal regions contain deep green blocks (similarity 0.6–0.8), indicating significant correlations in expert selection among different scaffold categories. Figure 3b, after introducing scaffold contrastive loss, shows a general reduction in off-diagonal similarity (0.2–0.4), demonstrating that scaffold contrastive loss function effectively promotes feature decoupling among scaffold-level experts, thereby enhancing the model’s specificity in learning.

**Figure 3:**
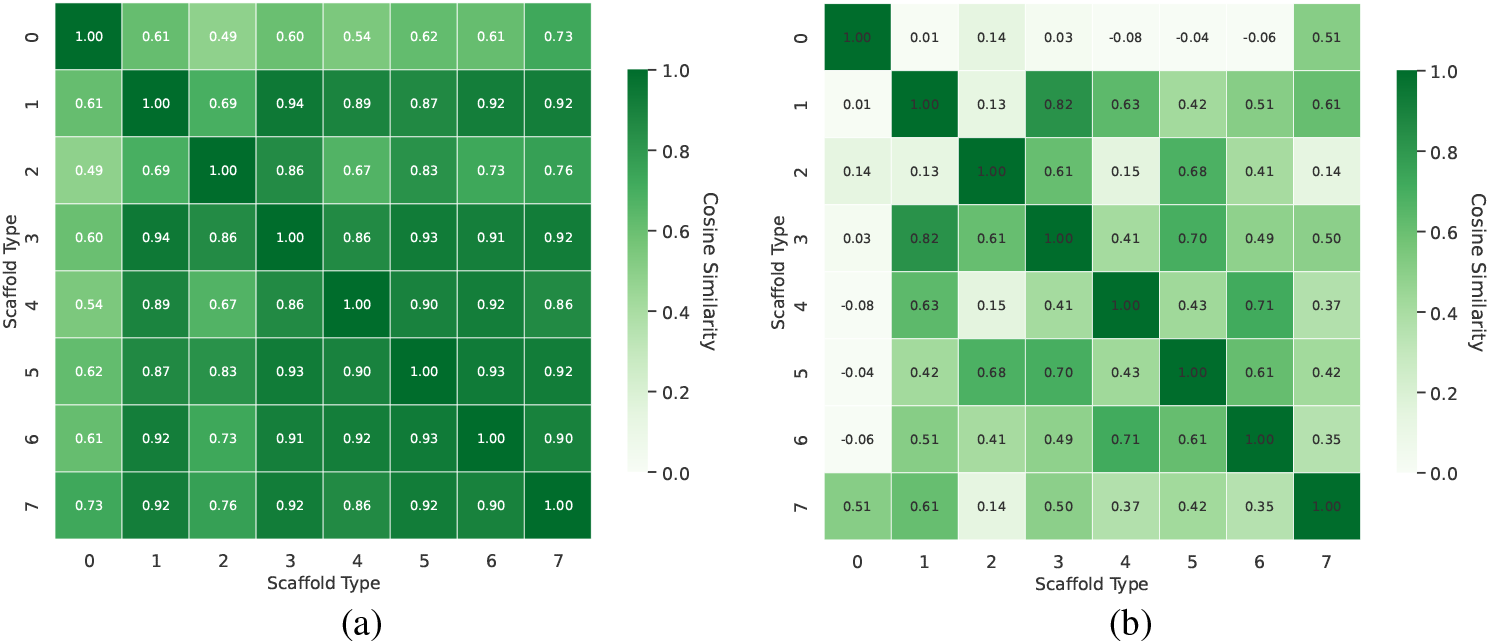
Comparison of expert selection distribution similarity (a) without and (b) with scaffold contrastive loss.

#### Importance of curriculum learning strategy

In Figure 4, we monitored the stability metric of atomic routing (Sim_*W*_ in Equation 9) throughout training and compared three different weighting methods. When using weight and mean strategies, dominant atoms in pretraining molecules (such as C and H) overwhelmingly influence training, leading to a stable Sim_*W*_ trend throughout training. However, when applying inverse weight, which prioritizes under-trained rare atoms, a noticeable routing shift emerges around epoch 10. Based on these observations, we integrate scaffold-assisted routing at the atomic-level routing stabilization phase (epoch >10) to prevent the model from learning in an incorrect direction.

**Figure 4:**
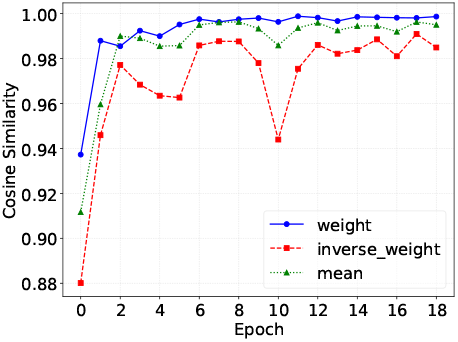
Trend analysis of expert selection similarity over training epochs.

#### Impact of expert balancing loss

Figure 5 compares sample distribution across eight experts (A–H) in H-MoE, with and without expert balancing loss. The results show that balancing loss reduces selection biases, leading to more uniform sample allocation (e.g., decreased selection frequency for expert F and increased for expert G). Figure 6 analyzes routing selection entropy during training, showing that with balancing loss, entropy stabilizes around 2.0—significantly higher than the 1.2–1.6 fluctuation range without it. Overall, balancing loss enhances expert selection stability and uniformity, improving model utilization.

**Figure 5:**
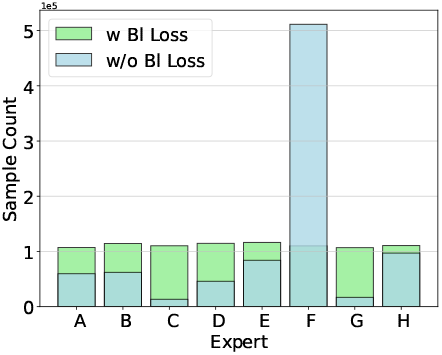
Impact of expert balancing loss on the final routing distribution of the model.

**Figure 6:**
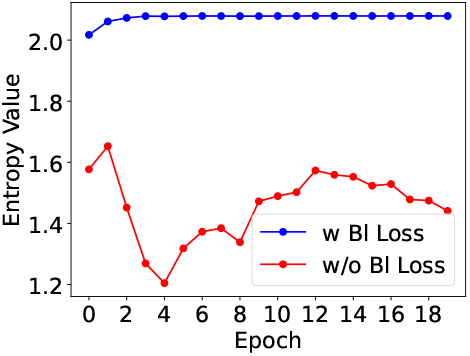
Impact of expert balancing loss on the entropy of routing choices during training.

#### Visualization and case study

In Appendix D.1, we present a visualization of scaffold-level and atom-level routing. Appendix D.2 provides t-SNE projections for different scaffold categories. Appendix D.3 examines the impact of various Morgan fingerprint bits on contrastive learning. Additionally, Appendix D.4 features a case study illustrating molecular scaffold structures across different categories.

## 6 Conlcusion

We propose a hierarchical scaffold-atom guided mixture-of-experts model for molecular representation learning. Theoretically, we demonstrate the effectiveness of scaffold-guided routing compared to traditional route strategies. Experimentally, our method, despite relying solely on 2D molecular information, surpasses some approaches that incorporate 3D conformations. Overall, we introduce a novel perspective to molecular representation learning, ensuring efficient modeling of diverse scaffold structures in pre-trained molecular datasets. Limitations are discussed in the Appendix E.

## A Theoretical analysis

### Theoretical motivation

We establish that scaffold-aware MoE models achieve provably better generalization under distribution shifts by: (A.1) decomposing molecular data distributions into scaffold-specific sub-domains, (A.2 and A.3) bounding cross-domain interference through gating error propagation, and (A.4) controlling capacity via scaffold-conditional Rademacher complexity. The results (A.5) indicate that under the condition that the minimum distance between different scaffolds is greater than zero, the scaffold-aware gating mechanism can significantly reduce the cross-domain interference term, thereby improving the model’s generalization performance.

### A.1 Problem formulation

#### Definition 1

(Scaffold-Partitioned Dataset). *Given molecular dataset* 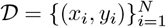 *with scaffolds s*(*x*_*i*_), *a scaffold split divides 𝒟into 𝒟*_*train*_, *𝒟*_*val*_, *𝒟*_*test*_ *such that:*

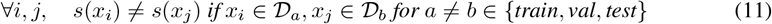

#### Definition 2

(MoE with Scaffold Awareness). *Our MoE model consists of:*

- *Experts:* 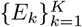 *where each E*_*k*_ *specializes on scaffold subset S*_*k*_
- *Gating network: g*(*x*) = [*g*_1_(*x*), …, *g*_*K*_(*x*)]^*T*^ *with*

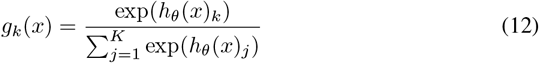

*where h*_*θ*_ : *X → ℝ*^*K*^ *is a scaffold-aware neural network*.

### A.2 Step 1: ideal expert specialization

#### Lemma 1

(Per-Expert Risk Decomposition). *For any expert E*_*k*_ *with ∥ E*_*k*_ *∥*_*∞*_ *≤M, the risk on test distribution decomposes as:*

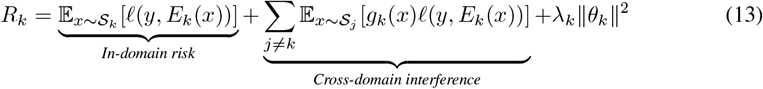

*where ℓ is an L-Lipschitz loss function and λ*_*k*_ *is the regularization parameter*.

*Proof*. Let *P*_*k*_ denote the distribution over *S*_*k*_. For any bounded expert *E*_*k*_ with parameters *θ*_*k*_:

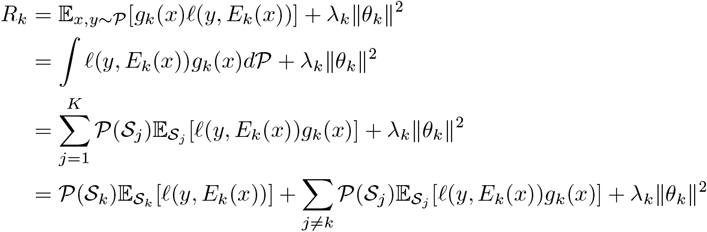

The inequality follows from the Lipschitz condition ∣ℓ (y, *E*_k_(x)) ∣≤ L∣y – *E*_k_ (x) ∣ ≤*L*(M +∥y∥_∞_).

### A.3 Step 2: gating network imperfection

#### Theorem 2

(Error Propagation Bound).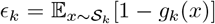 *be the expected gating error and* 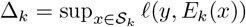. *Then with probability at least* 1 *™ δ:*

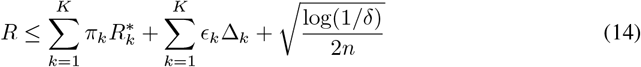

*where n is the sample size and π*_*k*_ = *P*(*S*_*k*_).

*Proof*. Using McDiarmid’s inequality, we first bound the deviation:

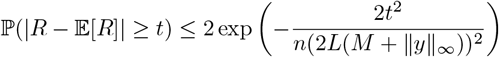

Setting the RHS to *δ* gives the concentration term. For the expectation:

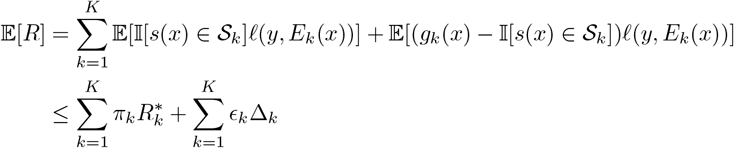

The result follows by combining these bounds via the union bound.

### A.4 Step 3: generalization guarantee

#### Theorem 3

(Scaffold-Aware Rademacher Complexity). *For K experts with* 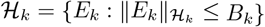 *and gating class 𝒢*= *{g* : *∥g∥*_*𝒢*_ *≤ B*_*g*_*}, we have:*

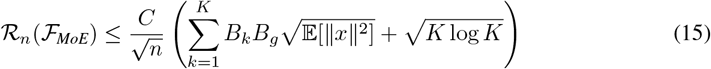

*where C is a universal constant*.

*Proof*. Using the duality between Rademacher complexity and covering numbers:

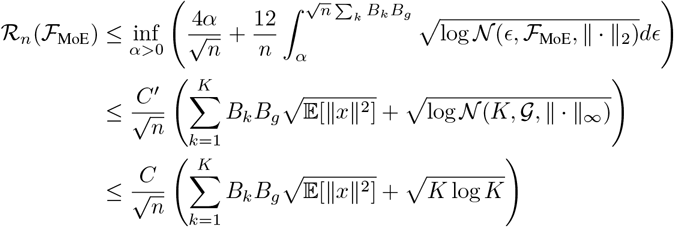

where we used the covering number bound *𝒩* (*ϵ, 𝒢,∥∥*_*∞*_) *≤* (3*B*_*g*_*/ϵ*)^*K*^ and Dudley’s entropy integral.

### A.5 Theoretical results

[Sample Complexity] For any *ϵ >* 0, to achieve *R ≤ R*^*∗*^ + *ϵ* with probability 1*™ δ*, the required sample size is:

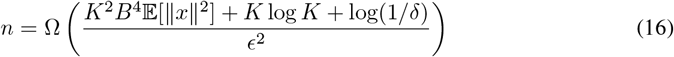

where *B* = max_*k*_ *B*_*k*_ *• B*_*g*_.

[Scaffold-Aware Benefit] Under the condition that min_*k≠ j*_ *d*(*S*_*k*_, *S*_*j*_) *≥ γ >* 0 where *d* is the Wasserstein distance, scaffold-aware gating reduces interference term by:

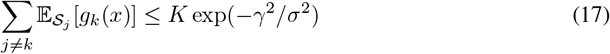

where *σ* is the Lipschitz constant of *h*_*θ*_.

## B Dynamic expert allocation

### Algorithm 2

Dynamic Expert Allocation

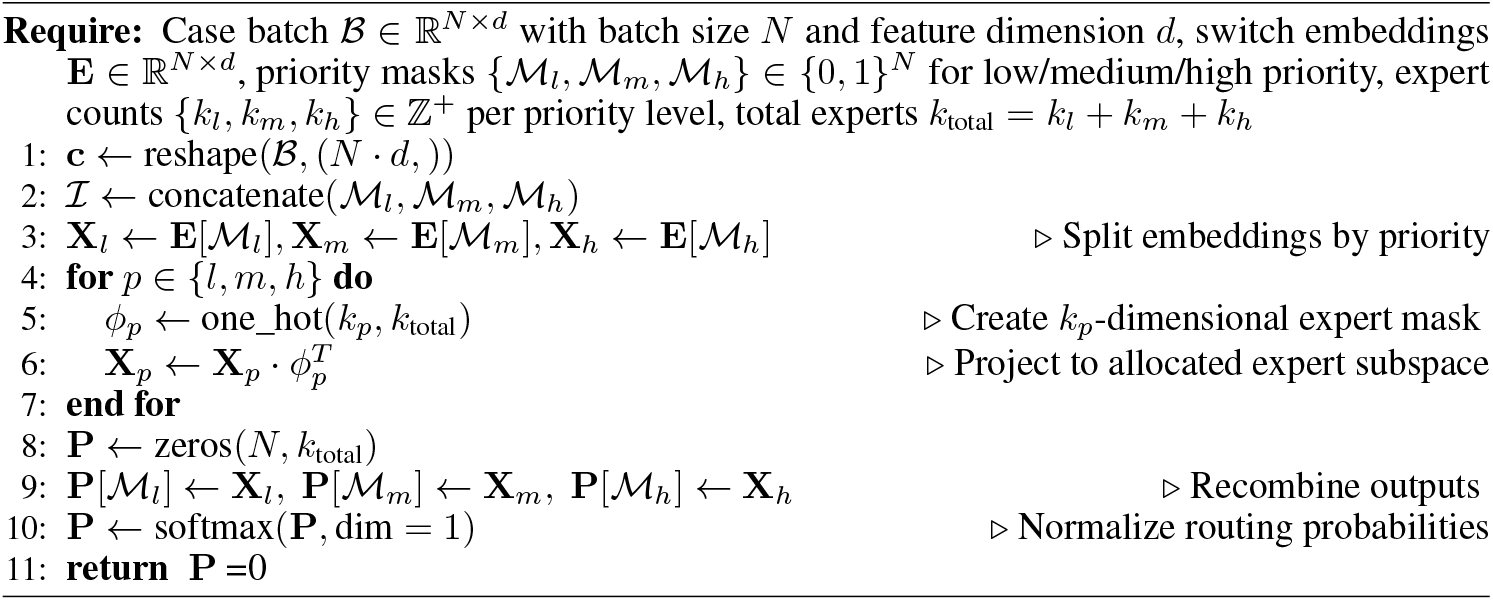

## C Experiment details

### C.1 Datasets details

MoleculeNet provides diverse datasets for molecular analysis across various applications.

- **HIV**. The HIV dataset contains 41,127 molecules screened for anti-HIV activity in a binary classification task.
- **BACE**. The BACE dataset, with 1,513 molecules, focuses on predicting BACE1 inhibition, relevant to Alzheimer’s drug discovery.
- **BBBP**. The BBBP dataset, comprising 2,039 molecules, investigates blood-brain barrier permeability using binary classification.
- **Tox21**. The Tox21 dataset, with 7,831 molecules, enables multi-label toxicity prediction, covering carcinogenicity and endocrine disruption.
- **SIDER**. The SIDER dataset, with 1,427 molecules, maps FDA-approved drugs to 27 types of side effects in a multi-label classification task.
- **ClinTox**. The ClinTox dataset, with 1,478 molecules, distinguishes approved drugs from those rejected due to toxicity.
- **MUV**. The MUV dataset, with 93,127 molecules, supports imbalanced binary classification for virtual screening reliability.
- **ToxCast**. The ToxCast dataset, with 8,576 molecules, extends high-throughput toxicity prediction across biological assays.
- **ESOL**. The ESOL dataset, with 1,127 molecules, predicts aqueous solubility (LogS), key for drug absorption.
- **FreeSolv**. The FreeSolv dataset, with 642 molecules, models solvation free energy in water, informing thermodynamic studies.
- **Lipo**. The Lipo dataset, with 4,200 molecules, assesses lipophilicity (LogD), influencing drug permeability.
- **Malaria**. The Malaria dataset, with 10,000 molecules, supports regression-based antimalarial screening via IC50 prediction.
- **CEP**. The CEP dataset, with 29,978 molecules, estimates photovoltaic efficiency, aiding organic solar cell research.

### C.2 Hyperparameters during pretraining and finetuning

We trained H-MoE on a single 24GB RTX 4090 GPU, completing pretraining within 24 hours. We have listed all the hyperparameters used during pretraining, as shown in the Table 4. For various downstream tasks, the hyperparameters we searched for include batch size, dense dropout, dropout rate, and learning rate. The hyperparameter combinations for each dataset are detailed in the Table 5.

**Table 4:** Hyperparameters for H-MoE in the pretraining stage.

| Hyperparameter | Value |
| --- | --- |
| Learning rate | 0.0001 |
| FFN dropout | 0.1 |
| Attention dropout | 0.1 |
| Embedding dropout | 0.1 |
| Encoder layers | 6 |
| Encoder attention heads | 4 |
| Encoder embedding dimension | 256 |
| Encoder FFN dimension | 512 |
| Vocabulary size | 15 |
| Activation function | GELU |
| Scaffold clusters | 16 |
| Initial expert number | 8 |
| Nbit of Morgan fingerprint | 2048 |
| Scaffold embedding dimension | 64 |

**Table 5:** Hyperparameters for various datasets.

| Dataset | Batch size | Dense dropout | Dropout rate | Learning rate |
| --- | --- | --- | --- | --- |
| ToxCast | 160 | 0.5 | 0.15 | 12e-5 |
| ClinTox | 64 | 0.1 | 0.05 | 14e-5 |
| Tox21 | 160 | 0.35 | 0.1 | 12e-5 |
| BBBP | 96 | 0.15 | 0.1 | 12e-5 |
| Sider | 64 | 0 | 0.2 | 8e-5 |
| HIV | 320 | 0.45 | 0.2 | 12e-5 |
| MUV | 480 | 0.2 | 0.15 | 10e-5 |
| Bace | 64 | 0.45 | 0.25 | 12e-5 |
| ESOL | 32 | 0.20 | 0.1 | 6e-5 |
| FreeSolv | 8 | 0.45 | 0.05 | 10e-5 |
| Lipo | 160 | 0.25 | 0.05 | 4e-5 |
| Malaria | 64 | 0.20 | 0.05 | 8e-5 |
| CEP | 960 | 0.15 | 0 | 8e-5 |

### C.3 Scaffold clusters in the pretraining and finetuning datasets

We have identified 16 distinct scaffold classes through k-means clustering from the ChEMBL dataset, which contains 1.45 million molecules. Their respective distributions are illustrated in Figure 7. The distribution of molecular scaffolds across clusters is highly imbalanced, with certain scaffold classes comprising the majority of samples. This is also the reason why we implement dynamic expert allocation strategies for different scaffolds categories. In the bar chart, we use different shades to show variations in sample quantity.

**Figure 7:**
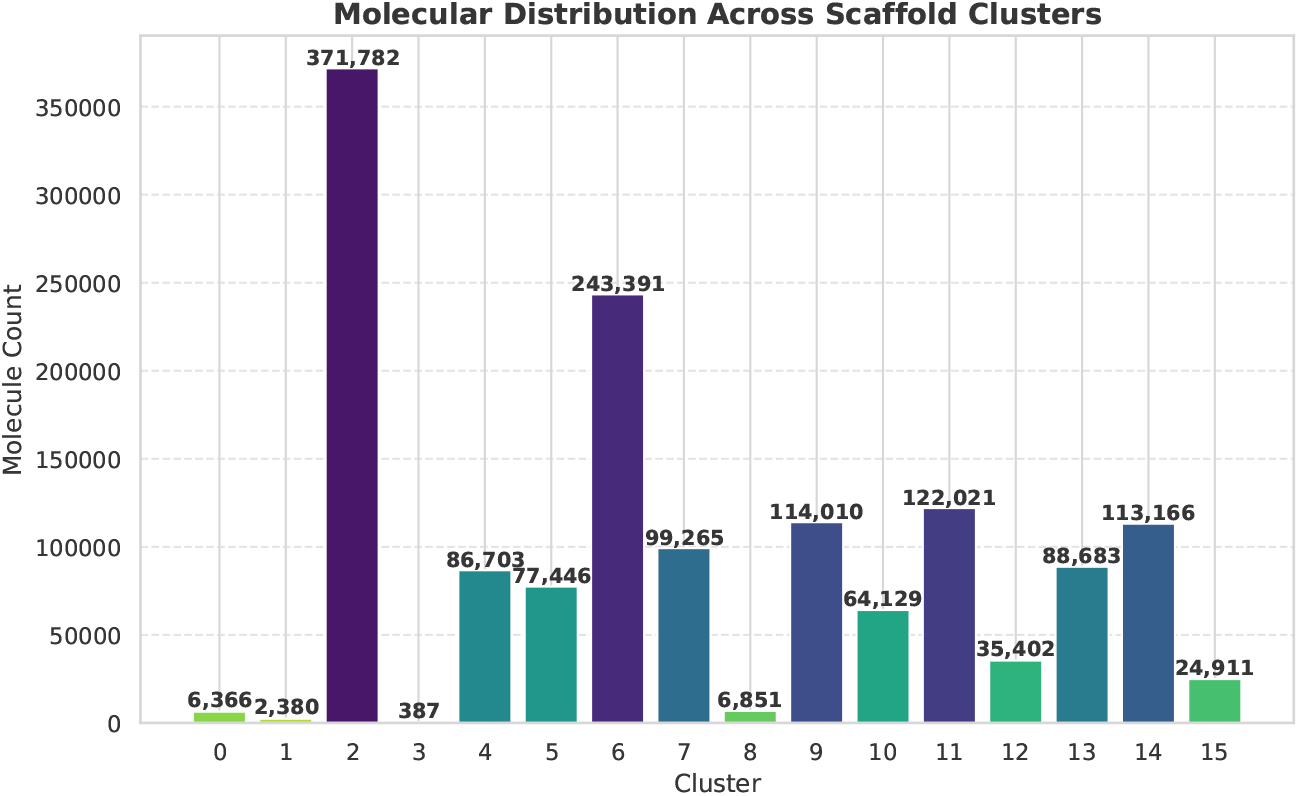
Molecule distribution across scaffold clusters in the pretraining dataset.

Similarly, in Table 6, we further present the scaffold distribution across eight classification datasets and five regression datasets. The scaffold composition in the downstream task datasets is also imbalanced. For instance, in FreeSolv, most molecules belong to Scaffold Class 5. Nevertheless, the large-scale pretraining-based scaffold-level routing strategy successfully captures similar patterns within individual scaffold clusters, enhancing the model’s ability to generalize across different scaffold distributions.

**Table 6:**
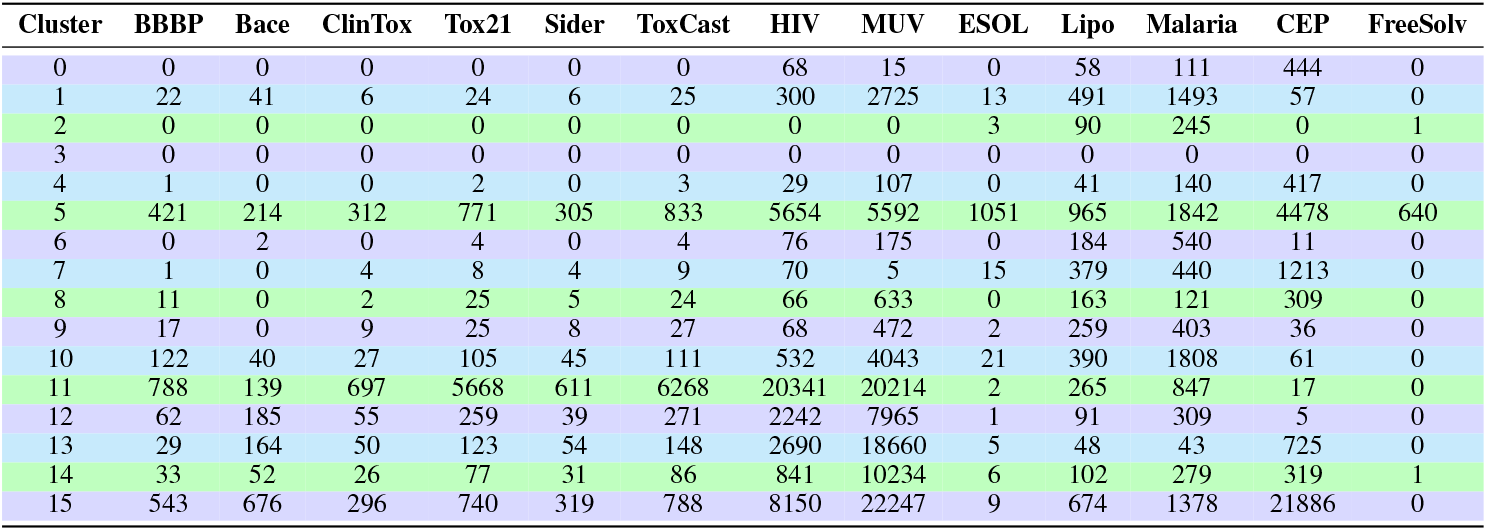
Molecule distribution across scaffold clusters in the downstream dataset.

## D Experiment results

### D.1 Visualization of routing distribution at atom and scaffold levels

Figures 8 and 9 visualize the routing distribution at both atomic and scaffold levels during pretraining (including experts discarded by the dynamic expert allocation strategy). The atomic-level routing distribution appears relatively uniform, primarily due to the expert balancing loss directly influencing the atomic-level router. Conversely, despite utilizing routing contrastive loss, scaffold-level routing remains concentrated, often assigning molecules to specific experts. This specialization allows certain experts to exclusively learn scaffold-level features. Overall, scaffold-level experts acquire specialized knowledge within distinct scaffold domains, while atomic-level experts capture fine-grained localized differentiations. This hierarchical global-local modeling strategy enhances the model’s capacity to process complex molecular features.

**Figure 8:**
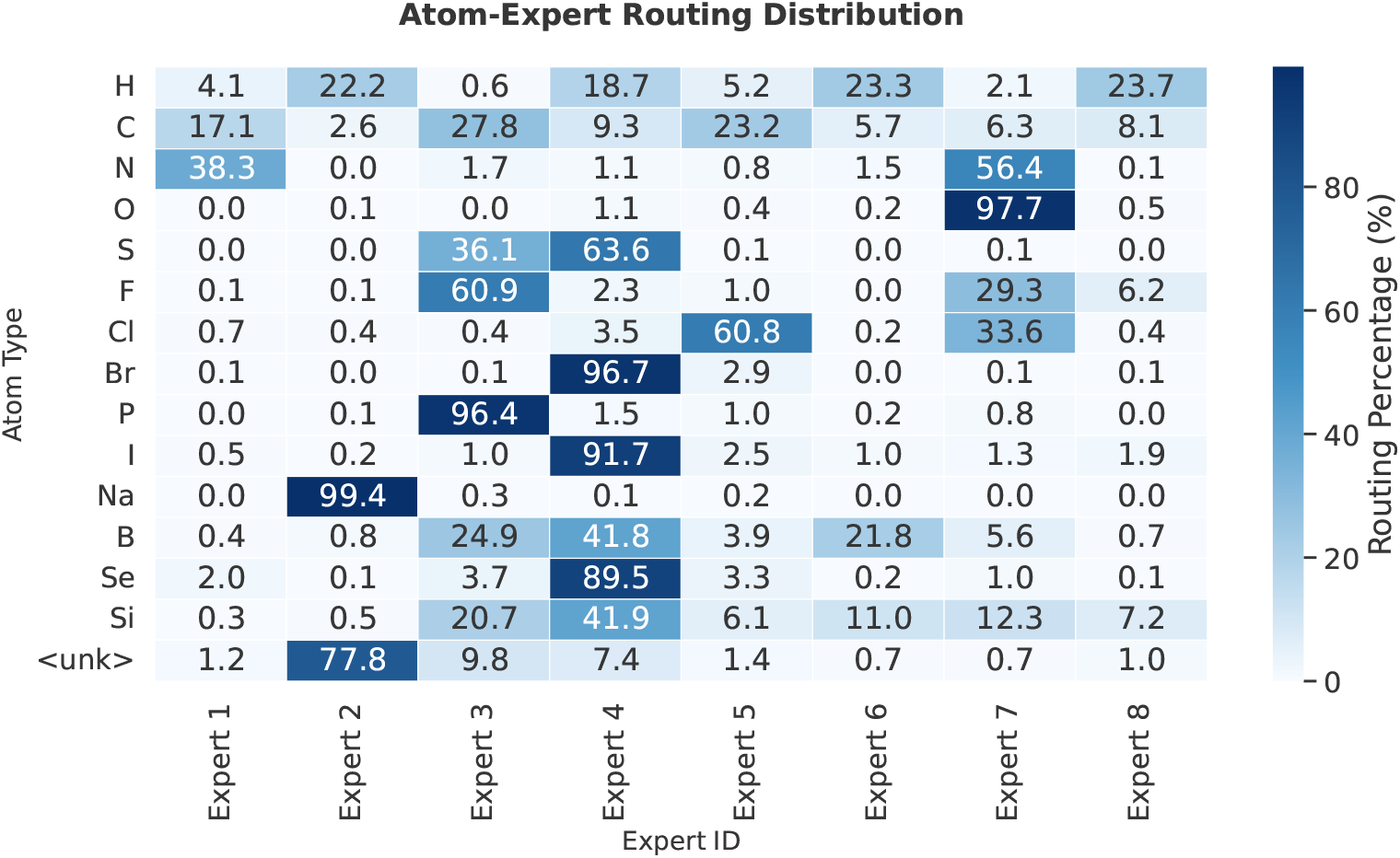
Atom-level expert routing heatmap of H-MoE.

**Figure 9:**
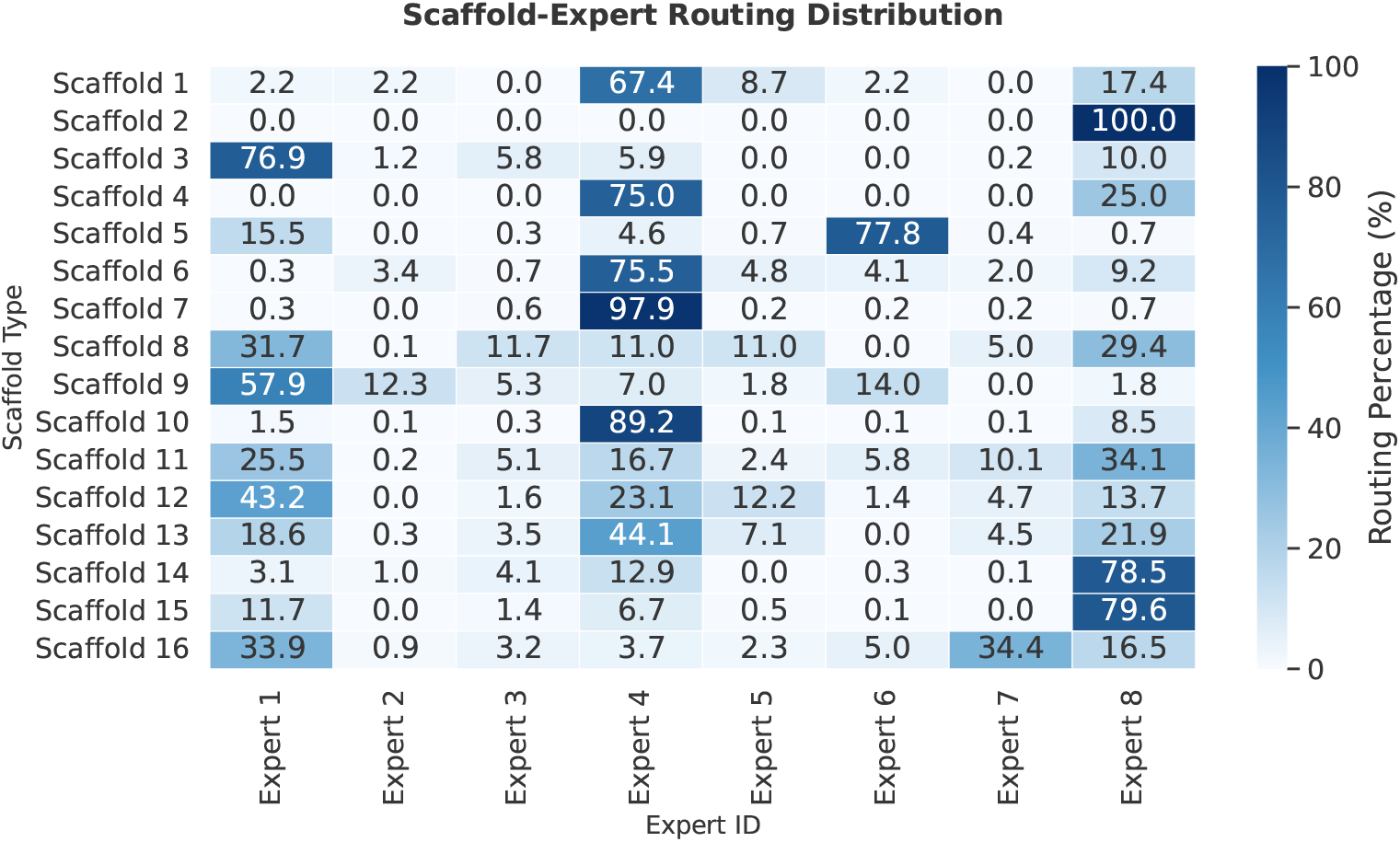
Scaffold-level expert routing heatmap of H-MoE.

Figure 10 visualizes the proportion of atomic types within each scaffold class in the pretraining dataset. Common atoms, such as hydrogen (H) and carbon (C), constitute a significant proportion, whereas other atomic types are relatively rare. However, some rare atoms play a crucial role in molecular properties, which is why inverse weighting is applied in Equation 9 to force the model to focus on these rare atoms. Additionally, Figure 10 illustrates how the hierarchical scaffold-atomic strategy enriches atomic categories, expanding the original 15 atomic types into 15 × 16 categories. This substantially enlarges the embedding vocabulary, enabling the model to learn more complex molecular features.

**Figure 10:**
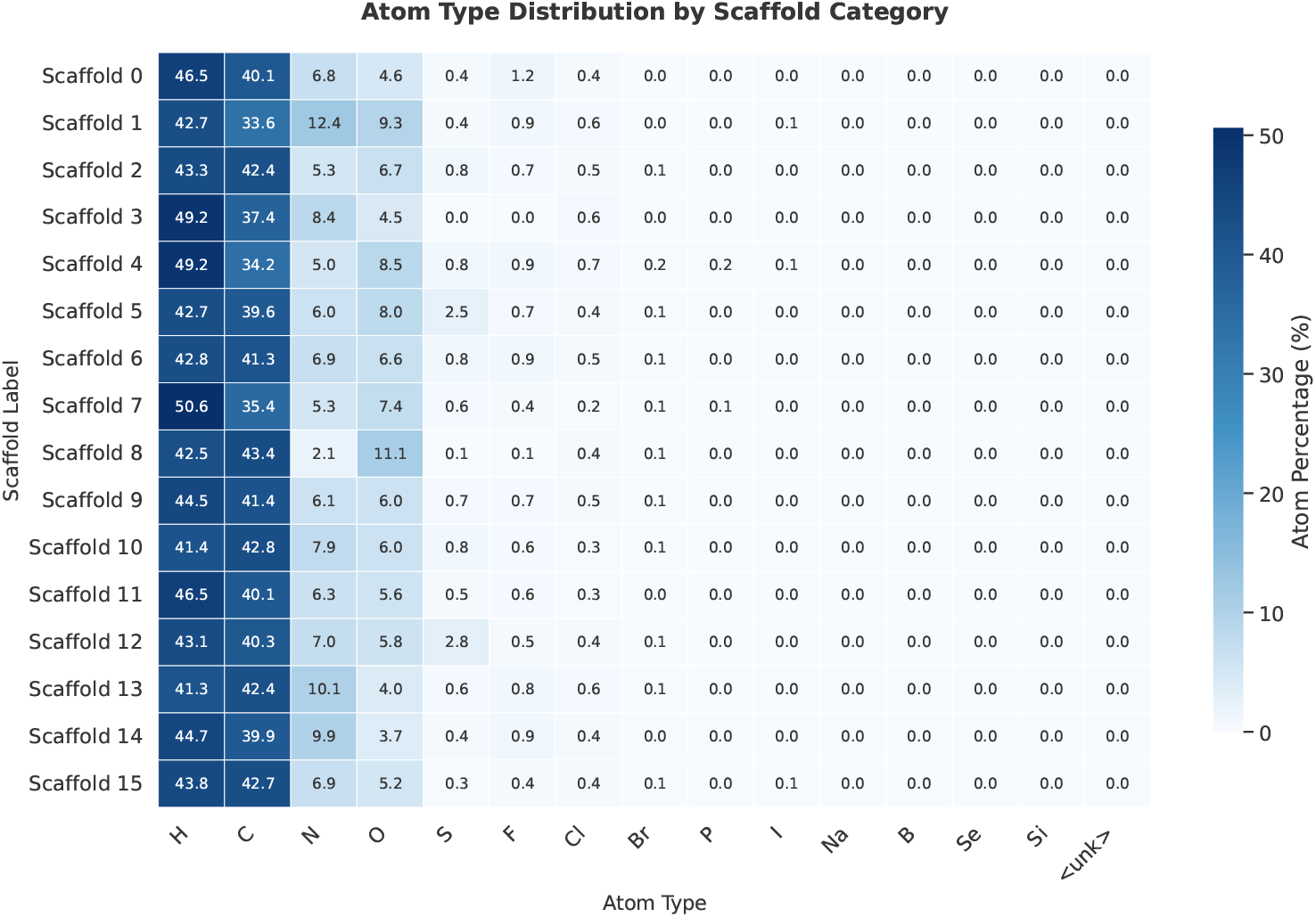
Atom type distribution by scaffold category.

### D.2 t-SNE plots of different scaffold classes

Figures 11–14 present the t-SNE dimensionality reduction plots of the embeddings for the top 4, top 8, top 12, and top 16 scaffold classes. Despite optimization using scaffold routing contrastive loss, as the number of scaffold classes increases, the boundaries between some rare scaffold clusters exhibit slight overlap. Given the overwhelming prevalence of common scaffolds compared to rare ones (as shown in Figure 7), the model is still able to learn general patterns within each scaffold category, thereby improving molecular representation capabilities.

**Figure 11:**
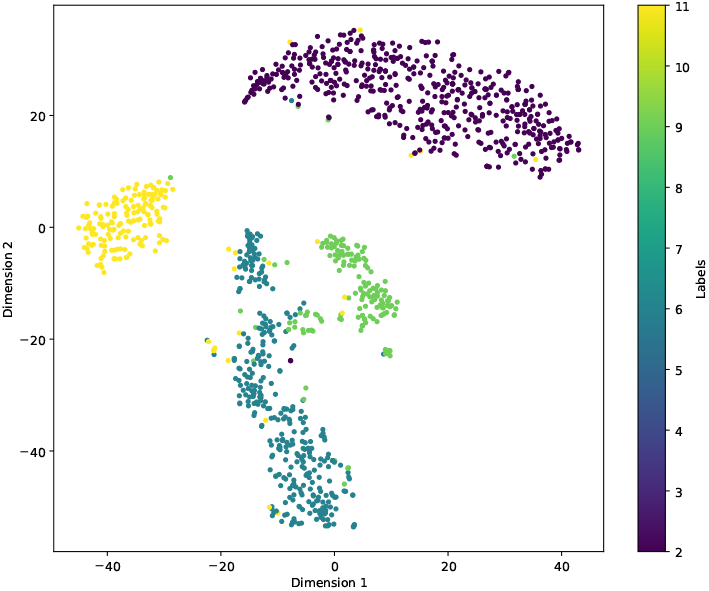
t-SNE plots of top-4 scaffold classes.

**Figure 12:**
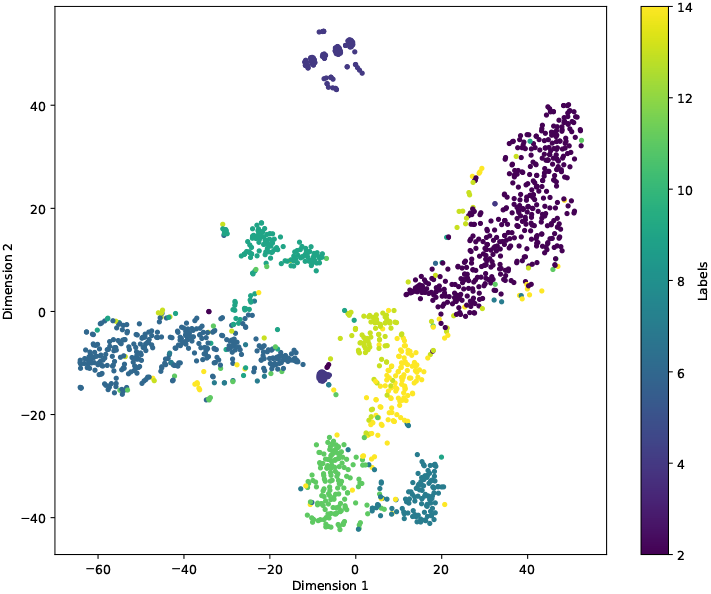
t-SNE plots of top-8 scaffold classes.

**Figure 13:**
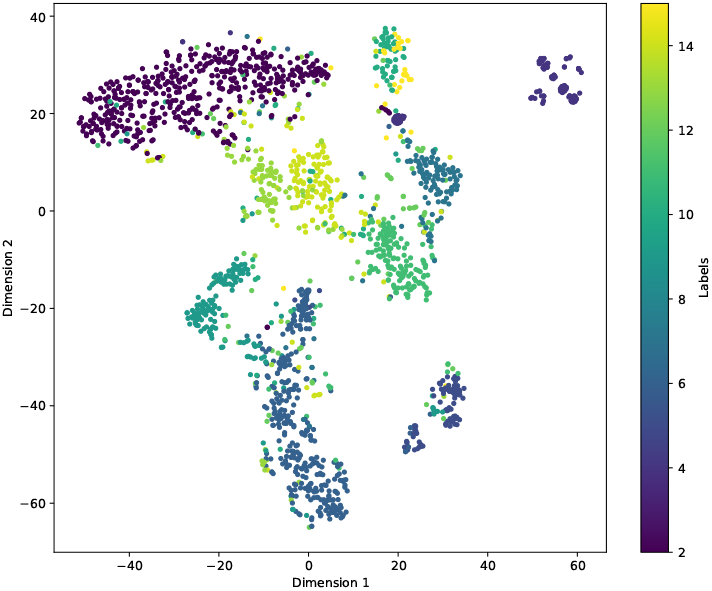
t-SNE plots of top-12 scaffold classes.

**Figure 14:**
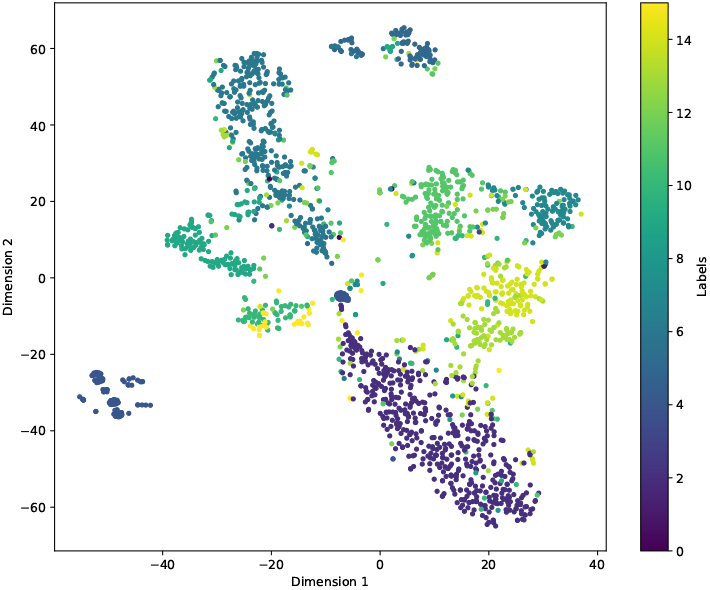
t-SNE plots of top-16 scaffold classes.

### D.3 Scaffold routing contrastive loss across different Morgan fingerprint bit

Figure 15 compares the convergence trends of molecular scaffold contrastive loss across different Morgan fingerprint bit lengths (nbit = 512/1024/2048/3072) over training epochs. The results indicate stable declines across all configurations, with larger fingerprint bit lengths achieving faster convergence. This is intuitive—higher bit lengths capture more intricate molecular structures. To balance computational efficiency and accuracy, we selected 2048 bits for the experimental setup.

**Figure 15:**
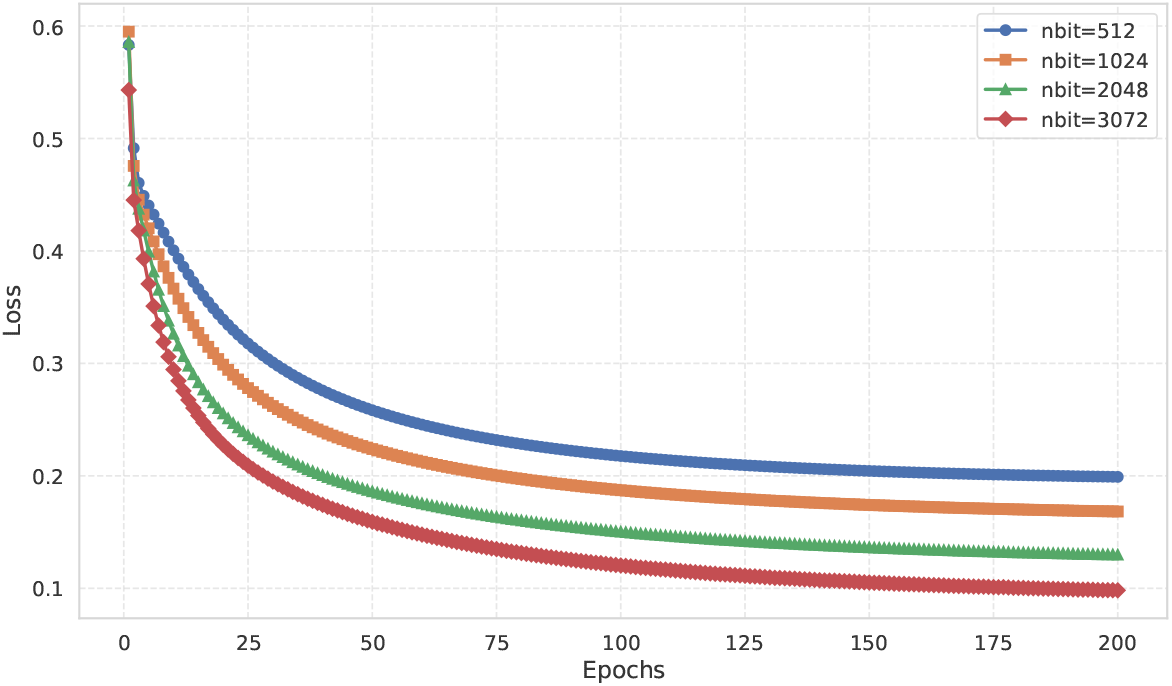
Scaffold routing contrastive loss across different Morgan fingerprint bit lengths over training epochs.

### D.4 Case study

Table 7 showcases the structural representations of six representative scaffolds (Scaffold 1–6) sampled from the pretraining dataset. Standardized chemical coloring (blue for nitrogen, red for oxygen, and yellow for sulfur) highlights key functional group differences. Each type of scaffold has relatively consistent features: Scaffold 1 displays aliphatic chains bridged by amide bonds between aromatic units; Scaffold 2 combines heterocyclic rings with amide groups, featuring nitrogen and carbonyl motifs; Scaffold 5 consists of fused aromatic systems decorated with amine and carbonyl functionalities. The topological differences among these scaffolds provide a structural basis for expert routing in subsequent tasks.

**Table 7:**
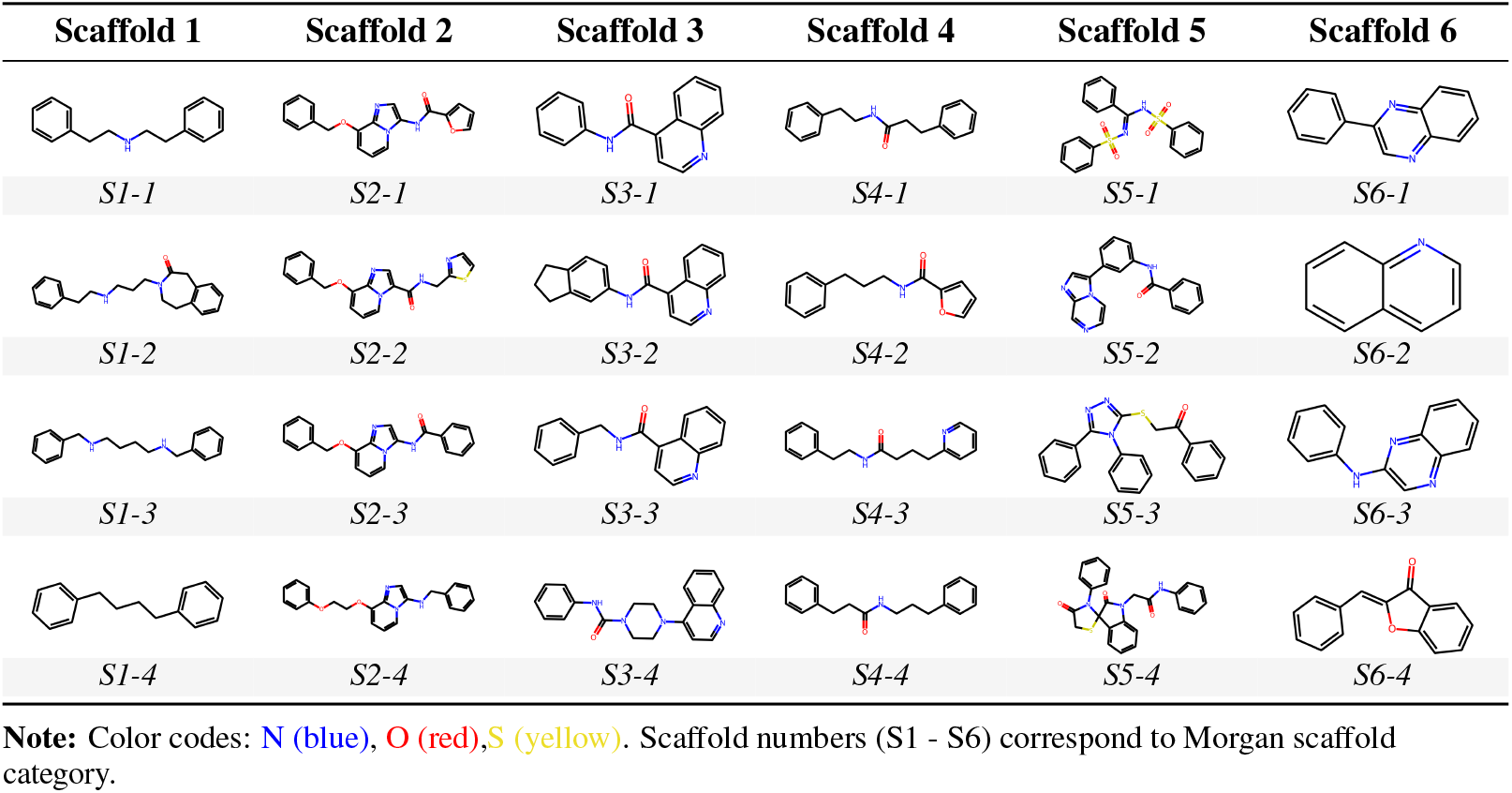
Visualization of different types of molecular scaffolds

## E Limitations

### Pretraining strategy

Our proposed H-MoE model currently employs the most fundamental masked pretraining task[33] within the transformer architecture. While this strategy has proven highly effective, we have observed other, more sophisticated pretraining approaches—such as MolBERT[9] and GraphMAE[7]—that may unlock even greater potential for H-MoE.

### 2D Graph vs. 3D conformation

Moreover, when processing molecular information, H-MoE has outperformed some methods[22][54] that incorporate three-dimensional (3D) conformation data, despite relying solely on two-dimensional (2D) features. This advantage can be attributed to several factors. **Firstly**, the 2D molecular information utilized in our method provides a more stable and generic representation, focusing on the topological structure and chemical bonds that are crucial for capturing the core properties of molecules. It avoids the noise and complexity introduced by the flexibility and diverse geometric configurations in 3D conformations. **Secondly**, our model’s innovative hierarchical scaffold-atom routing mechanism and contrastive loss function enable more effective capture of both global and local molecular features, further enhancing its ability to generalize across different molecular structures. **Additionally**, the dynamic expert allocation strategy ensures computational efficiency and adaptability to molecular diversity. These advantages collectively enable our model to achieve remarkable performance even in the absence of 3D molecular information.

However, from an information-theoretic perspective, 3D structural representations may contain more information than 2D graphs or 1D sequences, granting them a advantage in capturing the complexity of molecular structures. Therefore, exploring ways to integrate 3D feature information into H-MoE remains an intriguing and worthwhile topic for further discussion.

## References

[1] Meisheng Xiao, Qianhui Zheng, Paul Popa, Xinlei Mi, Jianhua Hu, Fei Zou, and Baiming Zou. Drug molecular representations for drug response predictions: a comprehensive investigation via machine learning methods. Scientific Reports, 15(1):20, 2025.

[2] Leo Breiman. Random forests. Machine learning, 45:5–32, 2001.

[3] Marti A. Hearst, Susan T Dumais, Edgar Osuna, John Platt, and Bernhard Scholkopf. Support vector machines. IEEE Intelligent Systems and their applications, 13(4):18–28, 1998.

[4] Jun Xia, Yanqiao Zhu, Yuanqi Du, and Stan Z Li. A systematic survey of chemical pre-trained models. arXiv preprint arXiv:2210.16484, 2022.

[5] Weihua Hu, Bowen Liu, Joseph Gomes, Marinka Zitnik, Percy Liang, Vijay Pande, and Jure Leskovec. Strategies for pre-training graph neural networks. arXiv preprint arXiv:1905.12265, 2019.

[6] Susheel Suresh, Pan Li, Cong Hao, and Jennifer Neville. Adversarial graph augmentation to improve graph contrastive learning. Advances in Neural Information Processing Systems, 34:15920–15933, 2021.

[7] Zhenyu Hou, Xiao Liu, Yukuo Cen, Yuxiao Dong, Hongxia Yang, Chunjie Wang, and Jie Tang. Graphmae: Self-supervised masked graph autoencoders. In Proceedings of the 28th ACM SIGKDD conference on knowledge discovery and data mining, pages 594–604, 2022.

[8] Chengxuan Ying, Tianle Cai, Shengjie Luo, Shuxin Zheng, Guolin Ke, D. He, Yanming Shen, and Tie-Yan Liu. Do transformers really perform badly for graph representation? Advances in neural information processing systems, 34:28877–28888, 2021.

[9] Jun Xia, Chengshuai Zhao, Bozhen Hu, Zhangyang Gao, Cheng Tan, Yue Liu, Siyuan Li, and Stan Z Li. Mole-bert: Rethinking pre-training graph neural networks for molecules. In The Eleventh International Conference on Learning Representations, 2023.

[10] Anna Gaulton, Louisa J Bellis, A Patricia Bento, Jon Chambers, Mark Davies, Anne Hersey, Yvonne Light, Shaun McGlinchey, David Michalovich, Bissan Al-Lazikani, et al. Chembl: a large-scale bioactivity database for drug discovery. Nucleic acids research, 40(D1):D1100–D1107, 2012.

[11] Zhenqin Wu, Bharath Ramsundar, Evan N Feinberg, Joseph Gomes, Caleb Geniesse, Aneesh S Pappu, Karl Leswing, and Vijay Pande. Moleculenet: a benchmark for molecular machine learning. Chemical science, 9(2):513–530, 2018.

[12] Jianyuan Deng, Zhibo Yang, Hehe Wang, Iwao Ojima, Dimitris Samaras, and Fusheng Wang. A systematic study of key elements underlying molecular property prediction. Nature Communications, 14(1):6395, 2023.

[13] Damai Dai, Chengqi Deng, Chenggang Zhao, RX Xu, Huazuo Gao, Deli Chen, Jiashi Li, Wangding Zeng, Xingkai Yu, Yu Wu, et al. Deepseekmoe: Towards ultimate expert specialization in mixture-of-experts language models. arXiv preprint arXiv:2401.06066, 2024.

[14] H Zhao, Z Qiu, H Wu, Z Wang, Z He, and J HyperMoE Fu. Towards better mixture of experts via transferring among experts. arXiv preprint cs.LG/2402.12656, 2024.

[15] Aran Komatsuzaki, Joan Puigcerver, James Lee-Thorp, Carlos Riquelme Ruiz, Basil Mustafa, Joshua Ainslie, Yi Tay, Mostafa Dehghani, and Neil Houlsby. Sparse upcycling: Training mixture-of-experts from dense checkpoints. arXiv preprint arXiv:2212.05055, 2022.

[16] Barret Zoph, Irwan Bello, Sameer Kumar, Nan Du, Yanping Huang, Jeff Dean, Noam Shazeer, and William Fedus. St-moe: Designing stable and transferable sparse expert models. arXiv preprint arXiv:2202.08906, 2022.

[17] Yuanhang Yang, Shiyi Qi, Wenchao Gu, Chaozheng Wang, Cuiyun Gao, and Zenglin Xu. Xmoe: Sparse models with fine-grained and adaptive expert selection. arXiv preprint arXiv:2403.18926, 2024.

[18] Xudong Lu, Qi Liu, Yuhui Xu, Aojun Zhou, Siyuan Huang, Bo Zhang, Junchi Yan, and Hongsheng Li. Not all experts are equal: Efficient expert pruning and skipping for mixture-of-experts large language models. arXiv preprint arXiv:2402.14800, 2024.

[19] Yanqi Zhou, Tao Lei, Hanxiao Liu, Nan Du, Yanping Huang, Vincent Zhao, Andrew M Dai, Quoc V Le, James Laudon, et al. Mixture-of-experts with expert choice routing. Advances in Neural Information Processing Systems, 35:7103–7114, 2022.

[20] Yuning You, Tianlong Chen, Yongduo Sui, Ting Chen, Zhangyang Wang, and Yang Shen. Graph contrastive learning with augmentations. Advances in neural information processing systems, 33:5812–5823, 2020.

[21] Shengchao Liu, Hanchen Wang, Weiyang Liu, Joan Lasenby, Hongyu Guo, and Jian Tang. Pre-training molecular graph representation with 3d geometry. arXiv preprint arXiv:2110.07728, 2021.

[22] Qiying Yu, Yudi Zhang, Yuyan Ni, Shikun Feng, Yanyan Lan, Hao Zhou, and Jingjing Liu. Multimodal molecular pretraining via modality blending. In The Twelfth International Conference on Learning Representations, 2024.

[23] Noam Shazeer, Azalia Mirhoseini, Krzysztof Maziarz, Andy Davis, Quoc Le, Geoffrey Hinton, and Jeff Dean. Outrageously large neural networks: The sparsely-gated mixture-of-experts layer. arXiv preprint arXiv:1701.06538, 2017.

[24] Dmitry Lepikhin, HyoukJoong Lee, Yuanzhong Xu, Dehao Chen, Orhan Firat, Yanping Huang, Maxim Krikun, Noam Shazeer, and Zhifeng Chen. Gshard: Scaling giant models with conditional computation and automatic sharding. arXiv preprint arXiv:2006.16668, 2020.

[25] William Fedus, Barret Zoph, and Noam Shazeer. Switch transformers: Scaling to trillion parameter models with simple and efficient sparsity. Journal of Machine Learning Research, 23(120):1–39, 2022.

[26] Joan Puigcerver, Carlos Riquelme, Basil Mustafa, and Neil Houlsby. From sparse to soft mixtures of experts. arXiv preprint arXiv:2308.00951, 2023.

[27] Xinyu Zhao, Xuxi Chen, Yu Cheng, and Tianlong Chen. Sparse moe with language guided routing for multilingual machine translation. In The Twelfth International Conference on Learning Representations, 2023.

[28] Quzhe Huang, Zhenwei An, Nan Zhuang, Mingxu Tao, Chen Zhang, Yang Jin, Kun Xu, Liwei Chen, Songfang Huang, and Yansong Feng. Harder tasks need more experts: Dynamic routing in moe models. arXiv preprint arXiv:2403.07652, 2024.

[29] Guy W Bemis and Mark A Murcko. The properties of known drugs. 1. molecular frameworks. Journal of medicinal chemistry, 39(15):2887–2893, 1996.

[30] Jian Zou, Zuo-Cheng Qiu, Qiang-Qiang Yu, Jia-Ming Wu, Yong-Heng Wang, Ke-Da Shi, Yi-Fang Li, Rong-Rong He, Ling Qin, Xin-Sheng Yao, et al. Discovery of a potent antiosteoporotic drug molecular scaffold derived from angelica sinensis and its bioinspired total synthesis. ACS Central Science, 10(3):628–636, 2024.

[31] Odin Zhang, Haitao Lin, Hui Zhang, Huifeng Zhao, Yufei Huang, Chang-Yu Hsieh, Peichen Pan, and Tingjun Hou. Deep lead optimization: Leveraging generative ai for structural modification. Journal of the American Chemical Society, 146(46):31357–31370, 2024.

[32] Kevin Yang, Kyle Swanson, Wengong Jin, Connor Coley, Philipp Eiden, Hua Gao, Angel Guzman-Perez, Timothy Hopper, Brian Kelley, Miriam Mathea, et al. Analyzing learned molecular representations for property prediction. Journal of chemical information and modeling, 59(8):3370–3388, 2019.

[33] Jacob Devlin, Ming-Wei Chang, Kenton Lee, and Kristina Toutanova. Bert: Pre-training of deep bidirectional transformers for language understanding. In Proceedings of the 2019 conference of the North American chapter of the association for computational linguistics: human language technologies, volume 1 (long and short papers), pages 4171–4186, 2019.

[34] Sebastian G Rohrer and Knut Baumann. Maximum unbiased validation (muv) data sets for virtual screening based on pubchem bioactivity data. Journal of chemical information and modeling, 49(2):169–184, 2009.

[35] Govindan Subramanian, Bharath Ramsundar, Vijay Pande, and Rajiah Aldrin Denny. Computational modeling of β-secretase 1 (bace-1) inhibitors using ligand based approaches. Journal of chemical information and modeling, 56(10):1936–1949, 2016.

[36] Ines Filipa Martins, Ana L Teixeira, Luis Pinheiro, and Andre O Falcao. A bayesian approach to in silico blood-brain barrier penetration modeling. Journal of chemical information and modeling, 52(6):1686–1697, 2012.

[37] Michael Kuhn, Ivica Letunic, Lars Juhl Jensen, and Peer Bork. The sider database of drugs and side effects. Nucleic acids research, 44(D1):D1075–D1079, 2016.

[38] Kaitlyn M Gayvert, Neel S Madhukar, and Olivier Elemento. A data-driven approach to predicting successes and failures of clinical trials. Cell chemical biology, 23(10):1294–1301, 2016.

[39] John S Delaney. Esol: estimating aqueous solubility directly from molecular structure. Journal of chemical information and computer sciences, 44(3):1000–1005, 2004.

[40] David L Mobley and J Peter Guthrie. Freesolv: a database of experimental and calculated hydration free energies, with input files. Journal of computer-aided molecular design, 28:711–720, 2014.

[41] David Mendez, Anna Gaulton, A Patrícia Bento, Jon Chambers, Marleen De Veij, Eloy Félix, María Paula Magariños, Juan F Mosquera, Prudence Mutowo, Michał Nowotka, et al. Chembl: towards direct deposition of bioassay data. Nucleic acids research, 47(D1):D930–D940, 2019.

[42] Francisco-Javier Gamo, Laura M Sanz, Jaume Vidal, Cristina De Cozar, Emilio Alvarez, Jose-Luis Lavandera, Dana E Vanderwall, Darren VS Green, Vinod Kumar, Samiul Hasan, et al. Thousands of chemical starting points for antimalarial lead identification. Nature, 465(7296):305–310, 2010.

[43] Johannes Hachmann, Roberto Olivares-Amaya, Sule Atahan-Evrenk, Carlos Amador-Bedolla, Roel S Sánchez-Carrera, Aryeh Gold-Parker, Leslie Vogt, Anna M Brockway, and Alán Aspuru-Guzik. The harvard clean energy project: large-scale computational screening and design of organic photovoltaics on the world community grid. The Journal of Physical Chemistry Letters, 2(17):2241–2251, 2011.

[44] Will Hamilton, Zhitao Ying, and Jure Leskovec. Inductive representation learning on large graphs. Advances in neural information processing systems, 30, 2017.

[45] Minghao Xu, Hang Wang, Bingbing Ni, Hongyu Guo, and Jian Tang. Self-supervised graph-level representation learning with local and global structure. In International conference on machine learning, pages 11548–11558. PMLR, 2021.

[46] Yu Rong, Yatao Bian, Tingyang Xu, Weiyang Xie, Ying Wei, Wenbing Huang, and Junzhou Huang. Self-supervised graph transformer on large-scale molecular data. Advances in neural information processing systems, 33:12559–12571, 2020.

[47] Yuning You, Tianlong Chen, Yang Shen, and Zhangyang Wang. Graph contrastive learning automated. In International conference on machine learning, pages 12121–12132. PMLR, 2021.

[48] Jun Xia, Lirong Wu, Jintao Chen, Bozhen Hu, and Stan Z Li. Simgrace: A simple framework for graph contrastive learning without data augmentation. In Proceedings of the ACM web conference 2022, pages 1070–1079, 2022.

[49] Zaixi Zhang, Qi Liu, Hao Wang, Chengqiang Lu, and Chee-Kong Lee. Motif-based graph self-supervised learning for molecular property prediction. Advances in Neural Information Processing Systems, 34:15870–15882, 2021.

[50] Chuang Liu, Yuyao Wang, Yibing Zhan, Xueqi Ma, Dapeng Tao, Jia Wu, and Wenbin Hu. Where to mask: structure-guided masking for graph masked autoencoders. In Proceedings of the Thirty-Third International Joint Conference on Artificial Intelligence, pages 2180–2188, 2024.

[51] Chuang Liu, Zelin Yao, Yibing Zhan, Xueqi Ma, Dapeng Tao, Jia Wu, Wenbin Hu, Shirui Pan, and Bo Du. Hi-gmae: Hierarchical graph masked autoencoders. arXiv preprint arXiv:2405.10642, 2024.

[52] Hannes Stärk, Dominique Beaini, Gabriele Corso, Prudencio Tossou, Christian Dallago, Stephan Günnemann, and Pietro Liò. 3d infomax improves gnns for molecular property prediction. In International Conference on Machine Learning, pages 20479–20502. PMLR, 2022.

[53] Gengmo Zhou, Zhifeng Gao, Qiankun Ding, Hang Zheng, Hongteng Xu, Zhewei Wei, Linfeng Zhang, and Guolin Ke. Uni-mol: A universal 3d molecular representation learning framework. In The Eleventh International Conference on Learning Representations, 2023.

[54] Shengchao Liu, Weitao Du, Zhi-Ming Ma, Hongyu Guo, and Jian Tang. A group symmetric stochastic differential equation model for molecule multi-modal pretraining. In International Conference on Machine Learning, pages 21497–21526. PMLR, 2023.

[55] Xiao-Chen Zhang, Cheng-Kun Wu, Zhi-Jiang Yang, Zhen-Xing Wu, Jia-Cai Yi, Chang-Yu Hsieh, Ting-Jun Hou, and Dong-Sheng Cao. Mg-bert: leveraging unsupervised atomic representation learning for molecular property prediction. Briefings in bioinformatics, 22(6):bbab152, 2021.

